# Nuclear Import Defects Drive Cell Cycle Dysregulation in Neurodegeneration

**DOI:** 10.1101/2025.01.28.635269

**Authors:** Jonathan Plessis-Belair, Taylor Russo, Markus Riessland, Roger B Sher

## Abstract

Neurodegenerative diseases (NDDs) and other age-related disorders have been classically defined by a set of key pathological hallmarks. Two of these hallmarks, cell cycle dysregulation (CCD) and nucleocytoplasmic transport (NCT) defects, have long been debated as being either causal or consequential in the pathology of accelerated aging. Specifically, aberrant cell cycle activation in post-mitotic neurons has been shown to trigger neuronal cell death pathways and cellular senescence. Additionally, NCT has been observed to be progressively dysregulated during aging and in neurodegeneration, where the increased subcellular redistribution of nuclear proteins such as TAR DNA-Binding Protein-43 (TDP43) to the cytoplasm is a primary driver of many NDDs. However, the functional significance of NCT defects as either a primary driver or consequence of pathology, and how the redistribution of cell cycle machinery contributes to neurodegeneration, remains unclear. Here, we describe that pharmacological inhibition of importin-β nuclear import is capable of perturbing cell cycle machinery both in mitotic neuronal cell lines and post-mitotic primary neurons *in vitro*. Our *Nemf*^R86S^ mouse model of motor neuron disease, characterized by nuclear import defects, further recapitulates the hallmarks of CCD in mitotic cell lines and in post-mitotic primary neurons *in vitro*, and in spinal motor neurons *in vivo*. The observed CCD is consistent with the transcriptional and phenotypical dysregulation observed in neuronal cell death and cellular senescence in NDDs. Together, this evidence suggests that impairment of nuclear import pathways resulting in CCD may be a common driver of pathology in neurodegeneration.

**Graphical Abstract:** Graphical Abstract:
Overview of Dysregulated Cell Cycle Mechanisms in Neuronal Cells. A nuclear import block drives cell cycle re-entry from G_0_, culminating in cell cycle arrest at G_1_/S. This cell cycle arrest is associated with activation of CKIs from the INK locus (p15, p16, p18, p19) and Cip/Kip (p21, p27) which act on specific CDK/Cyclin complexes. This activity is further associated with the G_1_/S downregulation E2F and stathmins, resulting in microtubule dysregulation and cell cycle arrest.

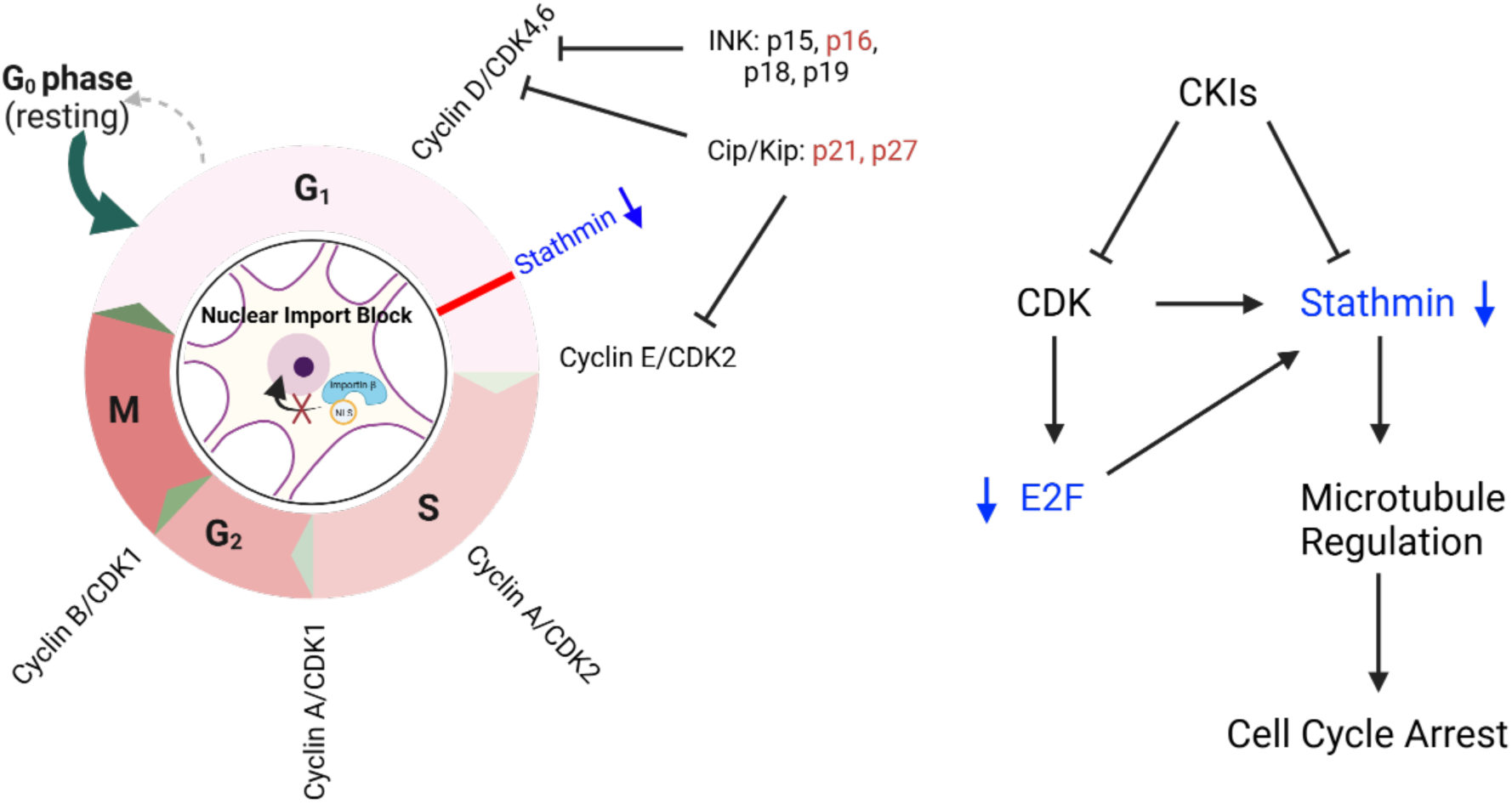

## Introduction

Aging is the primary risk factor for both cancer and NDDs (Hou et al., 2019; Sedrak & Cohen, 2023). Both replicative and physiological aging are intimately linked to cell-cycle decisions, and the ability of cells to undergo cell-cycle arrest in response to different triggers is a crucial process required for the maintenance of genomic integrity (Pietenpol & Stewart, 2002). Wherein CCD has traditionally been associated as a hallmark of tumor cells, the inappropriate activation of cell cycle regulators has also been implicated in the pathogenesis of NDDs (Herrup, 2012; Herrup & Arendt, 2002; Herrup & Yang, 2007; Martínez-Cué & Rueda, 2020; Stewart et al., 2003; Wang et al., 2009).

Proliferating eukaryotic cells that undergo regular cell divisions can be separated into discrete cell cycle phases: G_1_, S, G_2_, and M (Israels & Israels, 2000). The progression of cells from G_1_ to S and from G_2_ to M is regulated by restriction checkpoints (Blagosklonny & Pardee, 2002). These checkpoints serve to detect errors in the cell cycle prior to progression into the next phase and will drive cell-cycle arrest until the defect is repaired. Specifically, cyclins (Cyclin A, B, D, E) and cyclin-dependent kinases (CDK 1, 2, 4, 6) will form complexes and function to phosphorylate proteins which are crucial for cell cycle progression (Giacinti & Giordano, 2006). Phosphorylation of proteins such as retinoblastoma protein (RB) will result in the release of E2F transcription factors (TFs) and the activation of transcription of key genes that are required for progression through the G_1_/S restriction checkpoint (Möröy & Geisen, 2004; Siu et al., 2012; Topacio et al., 2019). Of these key genes, E2Fs will transcriptionally regulate stathmins for microtubule stability and cell cycle progression (Polager & Ginsberg, 2003; Polzin et al., 2004). The regulation of stathmins, specifically *STMN1* and *STMN2,* has been described to play an important role in motor neuron diseases and neurodegeneration (Baughn et al., 2023; Bellouze et al., 2016; Gagliardi et al., 2022; Klim et al., 2019; San Juan et al., 2022).

The regulation of cell cycle checkpoints is further associated with CDK inhibitors (CKIs) which can be broken down into two main families (Fischer et al., 2003). The Cip/Kip family includes the CKIs p21^Cip1^ (*CDKN1A*) and p27^Kip1^ (*CDKN1B*), which will actively inhibit CDKs 2, 4, and 6 (Pavletich, 1999). Therefore, p21 serves as a crucial regulator of cell cycle progression through the restriction checkpoint at the G_1_/S transition (Cayrol et al., 1998; Lepley & Pelling, 1997; Waldman et al., 1995). The second family of CKIs is dependent on the INK4 (Inhibitors of CDK4) gene locus whose four members p16^INK4a^ (*CDKN2A*), p15^INK4b^ (*CDKN2B*), p18^INK4c^ (*CDKN2C*), p19^INK4d^ (*CDKN2D*), exclusively bind and inhibit CDK4 and 6 (Roussel, 1999; Sherr & Roberts, 1999, 2004). Expression of this CKI family results in cell cycle arrest in the G_1_ phase and is accomplished by the inhibition of RB phosphorylation and repression of E2F TFs (Johnson et al., 1994; Nevins, 2001).

Balancing cell growth with division to maintain cellular homeostasis is a critical component of aging, whether cells are actively replicating or have entered a non-proliferative state. Neurons are considered to be post-mitotic and permanently arrested in the G_0_ phase of the cell cycle (Aranda-Anzaldo & Dent, 2017). However, neuronal cells that re-enter the cell cycle will often fail to complete the cell cycle and terminate in apoptosis or neurescence (neuronal cellular senescence) (Hudson et al., 2024; Jurk et al., 2012; Nandakumar et al., 2021; Riessland et al., 2019). In neurodegeneration, ectopic expression of cell cycle machinery and neuronal cell cycle re-entry is commonly observed as an early precursor of NDDs (Herrup & Arendt, 2002; Ruijtenberg & van den Heuvel, 2016; Yang et al., 2006). Currently, there is ambiguity on how these post-mitotic neurons re-enter the cell cycle and where they arrest, as well as with the pathways driving neuronal cell death versus neurescence. Yet, it is clear that aberrant activation of these cell-cycle mechanisms in post-mitotic neurons is a central process in neuronal aging and neurodegeneration.

A point of intersection in aging and NDDs is the decline in function of NCT, resulting in the intracellular redistribution of proteins and the accumulation of cytoplasmic aggregates such as TDP-43 (Kim & Taylor, 2017). We recently described a Nuclear Export Mediator Factor (NEMF) mouse model of neurodegeneration which presented with NCT defects, specifically defective importin-β nuclear import, as well as with transcriptional and phenotypical hallmarks of neurodegeneration (Plessis-Belair et al., 2024). Furthermore, the Drosophila ortholog of NEMF, Caliban, has been previously described to mediate the G_1_/S transition through E2F1 regulation of the cell cycle (Song et al., 2018). Here, we describe that an importin-β nuclear import block in both mitotic neuronal cell lines and post-mitotic primary neurons can dysregulate the cell cycle and result in cell cycle arrest. We further show that a G_1_/S cell cycle arrest is sufficient to cause the downregulation of stathmins, specifically *STMN2*. When this nuclear import block persists, a subpopulation of the cells will stochastically undergo apoptosis, while the surviving cells will transition into a senescence-like state. This CCD is further described in our *Nemf*^R86S^ mouse model through recapitulation of a G_1_/S cell-cycle arrest. This cell-cycle arrest is reiterated by aberrant expression of markers of cell-cycle dysregulation in both *Nemf*^R86S^ primary neuronal cultures and spinal motor neurons. We provide evidence that defective importin-β nuclear import drives CCD which culminates in a cascade of transcriptional and homeostatic alterations. Our results implicate age- and gene-driven dysfunction in NCT as a primary upstream mechanism driving neurodegeneration via CCD.

## Methods

### SK-N-MC Cell Culture

SK-N-MC cells (ATCC) were cultured according to the vendor’s protocol. In brief, cells were maintained in ATCC-formulated Eagle’s Minimum Essential Medium (10% FBS, 1% PenStrep) at 37ᵒC in 5% CO_2_. Reagents used indicated in Table S1.

### WT-NEMF and R86S-NEMF Mouse Embryonic Fibroblasts (MEFs) Derivation and Culture

MEFs were extracted and cultured as previously described(Plessis-Belair et al., 2024). In brief, WT NEMF and R86S-NEMF MEFs were maintained in Dulbecco’s Modified Eagle’s Medium (10% FBS, Glutamine 1X (Glutamax), 1% PenStrep) at 37ᵒC, 5% CO_2_ and passaged every two days with Trypsin-EDTA (0.05%). Reagents used indicated in Table S1.

### Importazole Treatment Time Series

Cells were treated with 20uM of importazole (Sigma) for 2, 12, 24, 48, 96 (4 days), and 168 (7 days) hours at 37ᵒC 5% CO_2_.

### Cell Cycle Inhibitor Treatments

Cells were treated with Nocodazole (500nM), Deferoxamine (10uM), L-Mimosine (100uM) for 48 hours at 37ᵒC 5% CO_2_.

### Fluorescence Associated Cell Sorting (FACS) Cell Cycle Analysis

Cells were washed one time with PBS (1X) and trypsinized and resuspended in media. Suspended cells were centrifuged for 5mins at 500 x g. The supernatant was removed, and cells were resuspended in 1mL of 70% Ethanol and incubated for at least 2 hours with inversion. Cells were washed once with PBS (1X). Cells were centrifuged again for 5mins at 500 x g, the supernatant was removed, and cells were resuspended in 500μl DAPI/PBS/triton X-100 (0.1%). Cells were incubated for 30mins in the dark. FACS sorting was adjusted for excitation at 340nm to 380nm and detection of DAPI for G_1_/S/G_2_ discrimination.

### Immunofluorescence Staining

As previously reported(Plessis-Belair et al., 2024), cells were chemically fixed in 4% paraformaldehyde in PBS(1X). Each well was then rinsed with PBS(1X). The cells were then permeabilized with 0.1% TritonX-100/PBS for 10mins at room temperature. The cells were then blocked for 1 hr at room temperature with 5% NGS/PBS/0.1% Tween-20. The cells were then incubated with the respective primary antibody in 5% NGS/0.1% Tween-20/PBS overnight at 4ᵒC. The primary antibodies used are listed in Table S1. The next day, the cells were rinsed 3 times for 5mins with 0.1% Tween-20/PBS(1X). The cells are then incubated with the respective secondary antibodies in 5% NGS/0.1% Tween-20/PBS for 2hrs in the dark at room temperature. The secondary antibodies used are listed in Table S1. The cells were then rinsed 3 times for 5mins with 0.1% Tween-20/PBS and stored temporarily with 200µL of PBS(1X) in the dark at 4ᵒC. Using sterile slides, 20-25µL of Prolong glass with NucBlue (Thermo Fisher) was added to the slide. The liquid was then aspirated from the well and using an SE 5 tissue curved forceps, the coverslips were gently picked up and then placed with the cell layer (top) down on the slide. The slide was then cured in the dark at 4ᵒC for 24hrs. Confocal images were taken by an Olympus FV1000 Laser Scanning Confocal. Laser settings (laser strength, gain, and offset) and magnification were maintained across treatment groups. Post-processing of images was performed by ImageJ and Cell Profiler as described below.

### RNA Seq of IPZ-treated SK-N-MC neuronal cells and *Nemf*^R86S^ MEFs

Concentrated RNA was sent for bulk RNAseq to Azenta. In brief, sample quality control and determination of concentration was performed using TapeStation Analysis by Azenta, followed by library preparation and sequencing. Computational analysis included in their standard data analysis package was used for data interpretation. *Nemf*^R86S^ MEFs RNA-seq dataset was previously published(Plessis-Belair et al., 2024).

### Reconstruction and Inference of a Transcriptional Regulatory Networks

The reconstruction of transcriptional regulatory networks was performed as described in the TNI pipeline in the RTN package in R with the list of known human transcription factors (TFs) obtained from Fletcher et al 2013. In brief, mutual information between TFs (1192 regulons) and all potential targets (14022 targets) is computed by removing non-significant associations through permutation analysis (nPermutations=1000) with correction for multiple hypothesis testing (tni.permutation()). Unstable interactions are then removed through bootstrap analysis (tni.bootstrap()), creating a reference regulatory network (778232 edges). Next, the ARACNe algorithm is applied which utilizes the direct processing inequality (DPI) theorem to enrich regulons by eliminating the weakest interactions between two TFs and a common target gene (tni.dpi.filter()). The resulting network formed is herein referred to as the transcriptional regulatory network (60412 edges). From here, one can retrieve individual regulons and their weighted interactions with target genes (tni.get()).

### Two-tailed Gene Set Enrichment Analysis

Two-tailed gene set enrichment analysis are performed as described in the TNA pipeline in the RTN package in R. In brief, a transcriptional regulator analysis is performed to assess the overlap between each regulon and significantly differentially expressed genes in the dataset (tna.mra()). One-tailed gene set enrichment analysis (GSEA1) assesses the interaction between a regulon with a ranked gene list generated from the differentially expressed genes. The regulons are then scored based on the association between the differentially expressed genes and the resulting response or phenotype. Two-tailed gene set enrichment analysis (GSEA2) separates the differentially expressed genes into positive and negative targets based on Pearson’s correlation between the regulon and the targets and then assesses the positive or negative association between the regulon and gene targets. To evaluate this phenotype, a differential enrichment score (dES) is calculated based on a stepwise evaluation of positive (ES*pos*) and negative(ES*neg*) gene enrichment scores. A positive dES represents an activated regulon, whereas a negative dES represents a repressed regulon activity.

### Mitotracker and Lysotracker Visualization

For imaging of mitochondria and lysosomes, cells were plated on sterile 12 mm round glass coverslips and exposed to 7-day treatment with either DMSO or 20 μM IPZ. Following treatment, cells were washed with PBS 1X and then incubated for 15 minutes at 37°C in either 500 nM Mitotracker Red CMXRos in PBS 1X or 1 µM Lysotracker Deep Red in PBS 1X. Following treatment, cells were chemically fixed in 4% PFA and immunostained for other markers as described in *Immunofluorescence Staining*. Confocal images were taken by an Olympus FV3000 Laser Scanning Confocal. Laser settings (laser strength, gain, and offset) and magnification were maintained across treatment groups. Post-processing of images was performed by ImageJ and Cell Profiler as described below.

### Western Blotting

Western blotting was performed as previously described(Plessis-Belair et al., 2024). In brief, cells were lysed with RIPA lysis buffer (Sigma) supplemented with protease inhibitor. Protein concentrations were standardized by Pierce BCA Protein Assay. 10µg of lysate was prepared with 50mM dTT (BioRad) and 4X laemli buffer (BioRad). The lysates were then loaded onto a Stain-Free mini-protean 10 well pre-cast gel (BioRad) and mini-protean tank (BioRad) with a Chameleon 800 MW ladder (Licor). The gels were run at 200V for approximately 45mins in Tris/Glycine/SDS Running Buffer (BioRad). Proteins separated in gels were transferred to a PVDF membrane using a semi-dry blotting method. Transfers were run for 90mins, with the current maintained between 80mA-240mA. The membrane is then incubated with TBS-Based Odyssey blocking buffer for 1hr at room temperature. The membrane is then incubated in blocking buffer supplemented with 0.1% Tween-20 and primary antibody overnight. The primary antibodies used are listed in the Table S1. The membrane is then washed with TBS-T (0.1% Tween-20) 3 times for 5mins. The membrane is then incubated with blocking buffer (0.1% Tween-20) and the respective secondary antibody at room temperature for 2hrs. The secondary antibodies used are listed in the Table S1. The membrane is then washed with TBS-T 3 times for 5mins. The membrane is imaged with the Odyssey Scanner (Licor).

### Antibody Array

Antibody array was prepared as described by Cell Cycle Phospho Array protocol by Full Moon Biosystems. In brief, proteins were extracted through RIPA Lysis as described above in Western Blotting. The resulting cell lysate was then purified through buffer exchange. The purified protein was then biotinylated using a Biotin/DMF solution (10mg/mL). The antibody array slide is then blocked, rinsed with distilled water, and then the biotinylated protein is coupled onto the slide. The slide is briefly washed, and the proteins are detected through Cy3-streptavidin. The slide is washed with wash buffer and then extensively rinsed with distilled water. The antibody array is then dried and scanned using a microarray scanner.

### Mouse strains, husbandry, and genotyping

All mouse husbandry and procedures were reviewed and approved by the Institutional Animal Care and Use Committee at Stony Brook University and were carried out according to the NIH Guide for Care and Use of Laboratory Animals. Tail tissue was lysed in proteinase K at 55ᵒC overnight and extracted DNA was used for genotyping. Genotyping for B6J-*Nemf*^R86S^ was performed via PCR using the following primers: forward primer specific to wild-type allele: 5′-AACATTTGAAGAGTCGGGGA-3′; forward primer specific to mutant allele: 5′-AACATTTGAAGAGTCGGGGT-3′; reverse primer common for both alleles: 5′-GCAGGTGGATGGTAGCAACG-3′.

### Primary Neuronal Cell Extraction and Culture

Primary neuronal cell extraction was performed as previously described(Plessis-Belair et al., 2024). In brief, brains were quickly dissected from P0/P1 pups in 2mL of Hibernate-A/B27(0.5mM GlutaMAX, PenStrep, 1% B27). Cortices were minced and digested in Papain Digestion Medium (100units Papain in 2mL Hibernate-A). Slices were washed twice in 2mL of Hibernate-A/B27. Slices were then triturated 10 times with a siliconized 9-inch Pasteur pipette with a tip fire polished to an opening of 0.7 to 0.9µm diameter. Supernatants were then transferred to a new tube and the cells were pelleted by centrifugation at 80 x g for 5mins. Cells were then counted by hemocytometer. The remaining cell pellet was resuspended in an appropriate volume of Neurobasal-A/B27(B27, 1X, PenStrep, 1X GlutaMax) and the cells were plated at 80% of plating volume. Neurobasal media was changed after 45mins of culture, and then was subsequently changed every two days.

### Spinal Cord Immunostaining

Spinal cords were immunostained as previously described(Plessis-Belair et al., 2024). In brief, spinal cords were surgically removed and chemically fixed in 4% PFA in PBS 1X. Spinal cords were placed in optimal cutting temperature (OCT) and stored in -80ᵒC. Spinal cords were cryosectioned at -20ᵒC into 30µm sections and then stored in PBS 1X at 4℃. Sections were permeabilized in 0.3% TritonX-100/PBS and blocked in 5% NGS/PBS/0.1% TritonX-100. Sections were then incubated with the respective primary antibody in 5% NGS/PBS/0.1% TritonX-100 overnight at 4ᵒC. The primary antibodies used are listed in Table S1. The spinal cords were then incubated with the respective secondary antibodies in 5% NGS/PBS/0.1% TritonX-100 for 2hrs in the dark at room temperature. The secondary antibodies used are listed in Table S1. Confocal images were taken by an Olympus FV1000 Laser Scanning Confocal. Laser settings (laser strength, gain, and offset) and magnification were maintained across treatment groups. Post-processing of images was performed by ImageJ and Cell Profiler as described below.

### qPCR

RNA (200ng) was reverse transcribed (Superscript IV Reverse Transcriptase (Thermo Fisher)) and the output volume of 20µL was diluted in nuclease-free water to 40µL for a working concentration of 5ng/µL. Real time PCR was performed using SYBR Green PCR Master Mix (Applied Biosystems) on a QuantStudio 3 System (Applied Biosystems) with reaction specificity confirmed by melt curve analysis. All comparisons (Control vs Experimental) for each qPCR reaction were run on the same qPCR plate and were run in a triplicate. For qPCR primer sequence, see Table S5.

### Image Analyses

Images were analyzed in bulk through Cell profiler. Z-projections were taken from each image by maximum intensity and then separated by fluorophore. Nuclear/Cytoplasmic (N/C) Ratios were taken by comparing area of nucleus (DAPI) and area of the cytoplasm (Phalloidin or brightfield). Nuclear and Cytoplasmic Intensities were standardized to area.

## Results

### IPZ-treated mitotic neuronal cell lines demonstrate cell cycle dysregulation consistent with the downregulation of Stathmins

Previously, we described the effects of a transient nuclear import block using a small molecular antagonist of importin-β, importazole (IPZ), on an SK-N-MC neuronal cell line (Plessis-Belair et al., 2024). The observations of TDP-43 proteinopathies and transcriptional dysregulation of *Stmn2* in this cell line implicated TDP-43 regulatory dysfunction in these IPZ-treated cells. It has been reported that human *STMN2* expression is directly regulated by TDP43 by binding to a cryptic exon that is lacking in mouse *Stmn2 (Baughn et al., 2023)*. However, IPZ treatment in mouse cell lines where TDP-43 regulation of *Stmn2* is absent shows the same downregulation of *Stmn2*, highlighting that *STMN2* regulation is not TDP43-exclusive, and that there is a parallel pathway downstream of nuclear import which can culminate in the observed transcriptional dysregulation.

To elucidate which mechanisms are involved in this parallel pathway, we isolated RNA from 2, 12, 24, 48, 96, and 168 hour-IPZ-treated SK-N-MC cells (doubling time: 48hours) and performed bulk-RNA sequencing (Fig 1A). With time, there is a clear increase in the number of significant differentially expressed genes (DEGs, padj<0.05, log2FC>1|log2FC<-1), with 320 significant DEGs at 2H, 1442 DEGs at 48H, and 3972 DEGs at 168H. Gene Ontology (GO) analysis of significant DEGs at 48H highlights dysregulation of expressed genes in the regulation of cell proliferation (GO:0008283), cell division (GO:0051301), and aging (GO:0007568), specifically highlighting genes involved in the G_1_/S transition of mitotic cell cycle (GO:0000082) (Fig 1B,C). The significance of DEGs enriched in the G_1_/S transition pathways disappear at 96H but return at 168H (Fig 1C). However, the enrichment of DEGs observed in cell cycle arrest (GO:0007050) are observed at 96H and 168H (Fig 1C). Similarly, GO analysis reveals pathway enrichment in G_2_/M transition of the cell cycle (GO:0000086), cell proliferation, cell division, aging, and mitotic nuclear division (GO:0007067) at 168H (Fig 1C).

**Figure 1:**
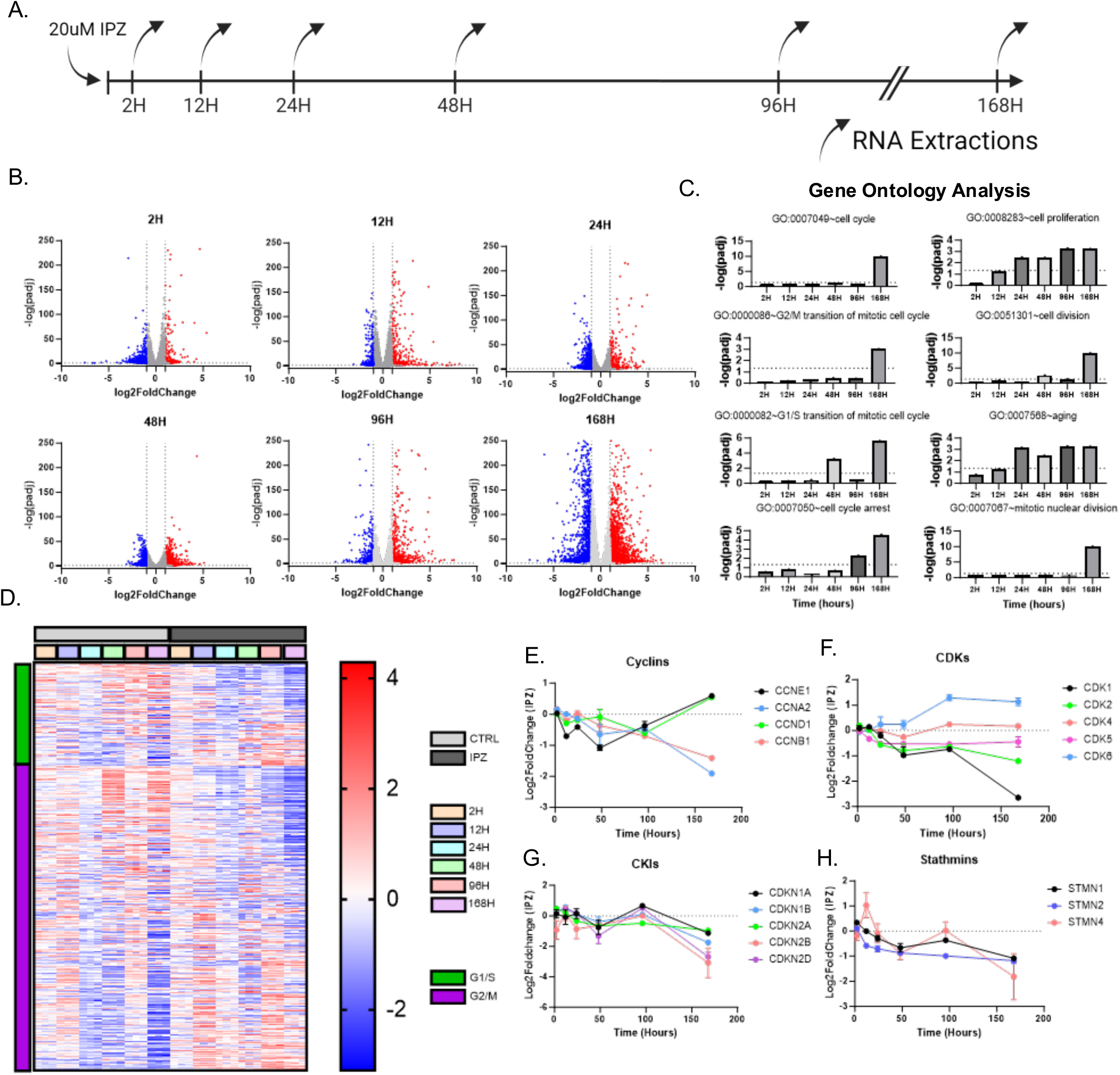
IPZ-treated mitotic neuronal cell lines demonstrate cell cycle dysregulation consistent with the downregulation of Stathmins. A) Timeline of the experimental set up. Cells were treated for 7 days in parallel with RNA extractions occurring at 2, 12, 24, 48, 96, and 168 hours. B) Volcano Plot of Significant DEGs for each time point. Red data points indicated upregulated DEGs (log2FC>1, padj<0.05) and blue data points indicate downregulated DEGs (log2FC<-1, padj<0.05). Gray data points indicate DEGs that do not meet the threshold (-1<log2FC<1|padj>0.05). C) Gene Ontology Analysis of significant DEGs shown in (B) for each given timepoint (padj<0.05). D) Z-score heatmap of genes associated with G_1_/S and G_2_/M clusters. E-H) Log2FoldChange Expression of Cyclins (E), CDKs (F), CKIs (G), Stathmins (H), over the 7-day time course.

To confirm whether these IPZ-treated cells were arrested at G_1_/S or G_2_/M, we utilized fluorescence-associated cell sorting (FACS) to isolate continuous DNA content profiles from IPZ-treated cells at each time point (Fig S1A)(Jayat & Ratinaud, 1993; Pozarowski & Darzynkiewicz, 2004). Control cells showed slight fluctuations in the percentage of cells in G_1_, S, and G_2_ throughout the time course, but did not show any overt cell cycle arrest as indicated by shifts into each peak (Fig S1B). IPZ-treated cells did not show any significant changes in the percentage of cells in G_1_, S, and G_2_ for the first 48 hours (Fig S1B). However, cells then accumulated into G_1_ peaks at 96H which was maintained until 168H (Fig S1B). This increase in G_1_ with a consistent decrease in S and no significant changes in G_2_ suggests a G_1_/S cell-cycle arrest initiated at around 96H (Fig S1B).

We then focused on genes associated with the G_1_/S and G_2_/M pathway and observed their differential expression over time (Fig 1D). Here, we observed phasic expression of cyclins, with their expression becoming increasingly dysregulated with time until we observed a split in expression, with G_1_/S associated *CCNE1* and *CCND1* being upregulated and G_2_/M associated *CCNB1* and *CCNA1* being downregulated at 168H (Fig 1E). Interestingly, expression of CDKs lack the phasic expression observed with cyclins, with *CDK6* showing an upregulation, and *CDK1* and *CDK2* showing a downregulation (Fig 1F). *CDK4* and *CDK5* show no greater dysregulation in expression (Fig 1F). Expression of CKIs similarly showed a more phasic cell-cycle expression profile, with all CKIs (*CDKN1A*, *CDKN2A*, *CDKN2B*, and *CDKN2D*) showing a general downregulation in expression at 168H (Fig 1G).

We sought to further investigate the dysregulation of cell cycle machinery by comparing the expression of long-noncoding RNAs (LNCRNAs) which have been implicated in both CCD and neurodegeneration(Hung et al., 2011; Kitagawa et al., 2013; Riva et al., 2016; Sun et al., 2015; Wan et al., 2017; Zhou et al., 2021). We analyzed significant DEGs of 5047 LNCRNAs which were further categorized into anti-sense, intronic, long interspersed noncoding (LINCRNA), long-noncoding non-systematic (LNC Non-systematic), microRNA, and small nucleolar RNA host genes (SNHG) (Fig S2A, B). Similar to the phasic expression of cell cycle components, LNCRNAs showed phasic expression, with alternating downregulation/upregulation of significant DEGs (Fig S2A, C). Specifically, LNCRNA *MEG3,* which has been implicated in neurodegeneration including Alzheimer’s Disease (AD), Parkinson’s Disease (PD), and Huntington’s Disease (HD) (Balusu et al., 2023; Chanda et al., 2018; Quan et al., 2020), showed an early upregulation at 48H with its expression returning to baseline by 168H (Fig S2D). In contrast, *MIR17HG* and *MIR22HG,* which have been implicated in AD and PD (Ning et al., 2022; Russo et al., 2024; Zhang et al., 2022), both show late upregulation at 168H with variable phasic expression throughout the time-course (Fig S2D).

We further investigated the transcriptional dysregulation of which have been shown to be de-repressed in neurodegeneration (Fondon et al., 2008; Li et al., 2012; Ravel-Godreuil et al., 2021; Reilly et al., 2013; Wojciechowska et al., 2014). We analyzed significant DEGs of 15295 simple repeat and TEs which were further categorized into DNA transposons, long interspersed nuclear elements (LINE), long terminal repeats (LTR), and short interspersed nuclear elements (SINE) (Fig S3A, B). Interestingly, we did not observe any phasic expression or dysregulation of simple repeats and TEs (Fig S3A, C). Rather, we saw a significant increase in dysregulated simple repeats and TEs at 168H, consistent with the large dysregulation observed with LNCRNA (Fig S3C). Altogether, the ultimate dysregulation of LNCRNA, simple repeats, and TEs suggests a general increase in heterochromatin relaxation consistent with aging and neurodegeneration. However, the phasic nature of dysregulation for LNCRNA suggests a more direct association with the observed cell-cycle dysregulation.

These alterations in cell cycle machinery were further associated with the dysregulation of stathmins, where *STMN1*, *STMN2*, and *STMN4* all showed a general downregulation with IPZ-treatment (Fig 1H). Interestingly, *STMN4* showed phasic expression similar to other cell cycle components, whereas *STMN2* showed a general downward trend in expression and *STMN1* fell somewhere in between (Fig 1H). Ultimately, inhibiting importin-β function resulted in cell-cycle dysregulation consistent with the cell-cycle associated downregulation of stathmins.

### IPZ treatment results in time-dependent cell-cycle regulator activity dysfunction

Given the critical role that the cell cycle plays in transcriptional regulation, we inferred and reconstructed a transcriptional network (RTN) of target genes and transcription factors (TFs) and subsequently performed two-tailed gene set enrichment analysis to isolate the activity of transcriptional regulators over time(Fletcher et al., 2013). Applying the ARACNe algorithm utilizing the data processing inequality (DPI) theorem to remove redundant and unstable interactions, we constructed an RTN comprised of 1192 regulons consisting of 14022 targets(see ‘Reconstruction and Inference of a Transcriptional Regulatory Networks’ in Methods). Regulon activity varied throughout the time course but remained balanced across samples throughout the RTN (Fig 2A).

**Figure 2:**
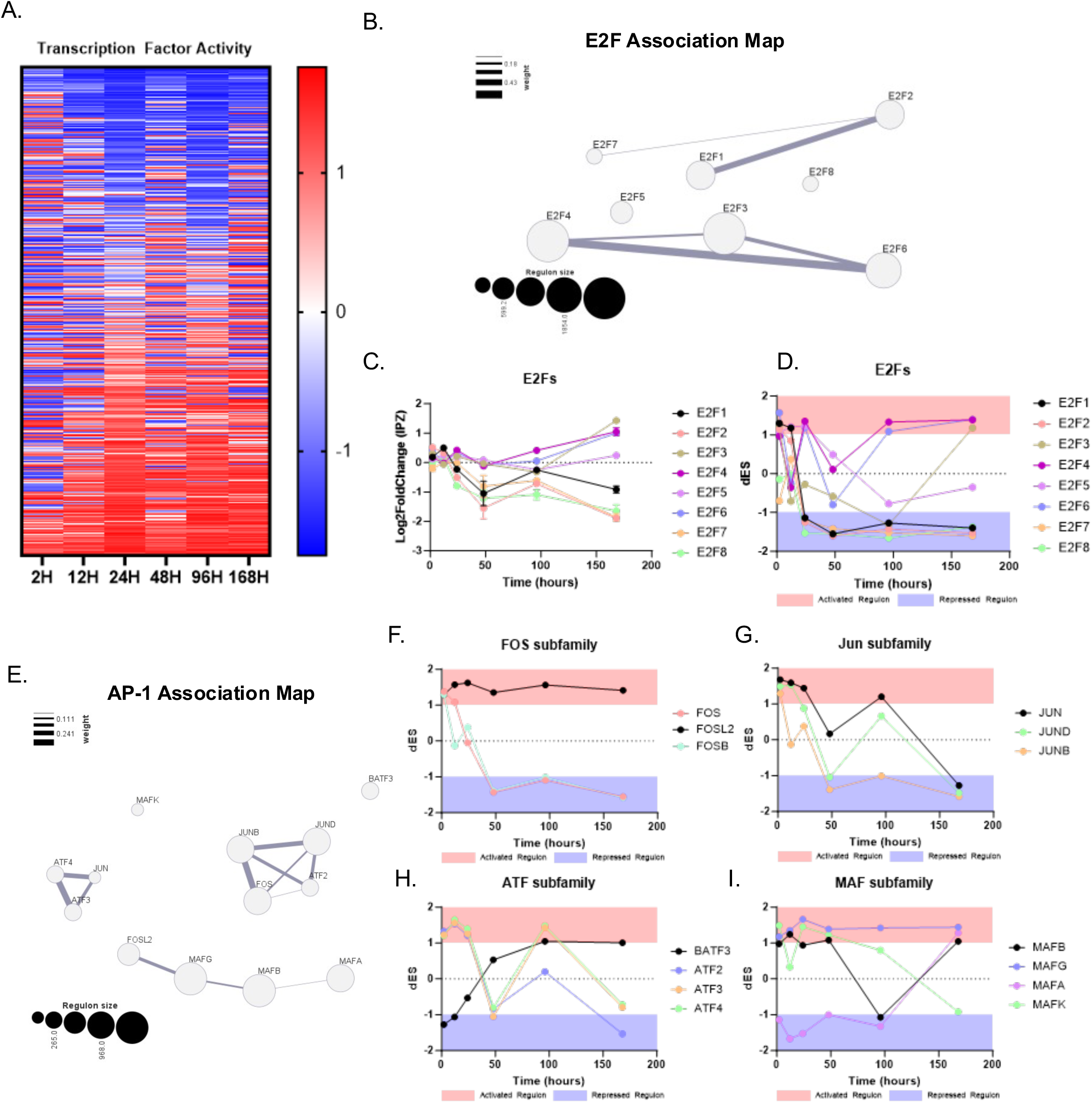
IPZ-treated cell lines show time-dependent cell-cycle regulator activity dysregulation. A) Differential Enrichment Score (dES) heatmap of IPZ-treated SK-N-MC cells. The activity of 1192 transcription factors are inferred based on a reconstructed transcriptional network (RTN) from SK-N-MC gene expression profiles. B) E2F Association Map inferred from the RTN displaying each regulon (E2Fs 1-8) and its size (represented by area), as well as overlapping associations in transcriptional activity with other regulons (measured by weighted line). C) Log2FoldChange Expression of E2Fs 1-8 over the 7-day time course. D) dES transcriptional activity of E2Fs 1-8 over the 7-day time course. E) AP-1 complex Association Map inferred from the RTN displaying each regulon family (FOS, JUN, ATF, and MAF). Size is represented by area as well as overlapping associations in transcriptional activity with other regulons (measured by weighted line). F-I) dES transcriptional activity of FOS (F), JUN (G), ATF (H), and MAF (I) subfamilies over the 7-day time course. Activated regulon activity in red shaded area (dES>1) and repressed regulon activity in blue shaded area (dES<-1). Scatter plot bars are mean with standard deviation (C).

From here, two-tailed gene set enrichment analysis revealed enrichment of transcriptional regulators in the E2F family. The E2F family of TFs has been described as key regulators for cell cycle progression through the G_1_/S checkpoint(Cam & Dynlacht, 2003; Zhu et al., 2004). We then reconstructed an association map and saw that E2F1 gene targets are linearly associated with E2F2 and E2F8, whereas E2F3 has a larger regulon size and greater association with E2F4 and E2F6 (Fig 2B). E2F5 and E2F8 are small regulons and do not show any preferential association in this RTN with other E2F TFs (Fig 2B). Observing the expression and activity of these E2F TFs revealed phasic expression profiles and activity of some TFs, but not all (Fig 2C,D). E2Fs 1, 2, 7, and 8 showed similar phasic expression patterns with a general downward trend in expression. Despite this phasic expression, these E2F TFs showed a significant repression in regulon activity as early as 24H which was maintained throughout the time course (Fig 2C). On the contrary, E2Fs 3, 4, 5, and 6 showed variable changes in expression correlated with an upregulation in expression and activity (Fig 2CD). Altogether, we suggest the E2F family of TFs as a primary upstream regulator for the observed cell-cycle associated transcriptional dysregulation.

AP-1 TFs have been shown to regulate the cell cycle by activating and inhibiting the expression of key components of cell-cycle machinery, including but not limited to *CCND1, CDKN1A, CDKN2B, CDKN2D,* and *TP53 (E. Shaulian & M. Karin, 2001; Shaulian & Karin, 2002)*. Thus, we turned to the transcriptional activity of factors associated with the AP-1 complex of transcriptional regulators. Here, we highlighted four subfamilies that can both hetero- and homo-dimerize to regulate transcriptional activity: Jun, FOS, MAF, and ATF(Bohmann et al., 1987; Chinenov & Kerppola, 2001; Fujiwara et al., 1993; Kataoka et al., 1995; Liebermann et al., 1998; Eitan Shaulian & Michael Karin, 2001). The construction of an AP-1 association map demonstrates that ATF4, ATF3, and JUN have high association and relatively small regulon sizes, whereas JUNB, JUND, FOS, and ATF2 show strong association of regulatory targets and slightly larger regulon sizes (Fig 2E). FOSL2, MAFG, MAFB, and MAFA are observed to be linearly associated, each with relatively large regulon sizes (Fig 2E). The activity of the FOS subfamily can be separated into FOS/FOSB which shows similar early and sustained repressed activity, whereas FOSL2 shows a maintained active regulon (Fig 2F). The Jun subfamily shows a more phasic activity pattern, with all three members JUN, JUND, and JUNB showing late repression of regulon activity (Fig 2G). The ATF subfamily consisting of ATFs 2, 3, and 4, showed similar phasic activity similar to the Jun subfamily, with the exception of BATF3 which becomes increasingly activated with time (Fig 2H). The MAF subfamily showed highly variable activity, with MAFA, B, and G showing a late activated regulon activity and MAFK showing a downward trend to repressed activity (Fig 2I).

Taken together, the time-dependent activity profiles provide insight on cell-cycle related activity changes versus general repression or activation of TFs as a result of the nuclear import inhibition, further highlighting nuclear import mediated cell-cycle dysfunction.

### Cell-cycle dysregulation is associated with hallmarks of senescence independent of CKI expression

The presence of CCD in the absence of CKI expression (*CDKN1A*, *CDKN2A/B*, *CDKN2D*, Fig 1G) even at our latest timepoints raises confounding questions on diverging pathways between apoptosis and senescence. Therefore, we aimed to explore whether surviving cells at 168 hours demonstrated canonical hallmarks of a senescence-like phenotype, including senescence-associated secretory phenotype (SASP), reduced lamin expression and increase in nuclear size, mitochondrial and lysosomal dysfunction, and DNA damage (Martínez-Cué & Rueda, 2020; Russo & Riessland, 2022).

Utilizing a previously established set of genes identified in senescence and the SASP (SenMayo)(Saul et al., 2022), we examined the expression of these genes over time following nuclear import inhibition and observed a significant dysregulation of genes associated with the SASP (Fig 3A). Specifically, *CXCL8* and *CCL20* upregulation occurred as early as 2 hours post-treatment, with a maintained expression increase throughout the time-course (Fig 3B). In comparison, *CXCL16*, *IL32*, *FGF2*, and *IL6ST* showed a time-dependent response correlated with a gradual increase in expression (Fig 3B). Interestingly, the most upregulated SASP factors lacked a phasic expression profile, suggesting that their upregulation was a cumulative response (Fig 3B). Examinations into nuclear envelope transcriptional regulation revealed a slight yet significant downregulation of lamins, specifically *LMNA* at 168H. The downregulation of *LMNB1* is small yet significant, wheras *LMNB2* showed less dysregulation (Fig 3C). Interestingly, the dysregulation of *LMNA* was strongly correlated with the phasic dysregulation of LINC complex component *SUN2*, but not *SUN1*, suggesting cell-cycle associated regulation of these gene transcripts which become increasingly more dysregulated with time (Fig 3C) (Haque et al., 2010). We then followed up on the dysregulation of lamins through immunostaining of senescent biomarker lamin B1, whose quantification showed a significant increase in nuclear size consistent with a decrease in lamin B1 intensity (Fig 3D-F). Qualitatively, the lamin B1 immunostaining displayed a subpopulation of cells with large, flat nuclei, with very few folds in the nuclear envelope, which is consistent with a senesence-like phenotype (Fig 3D) (González-Gualda et al., 2021; Neurohr et al., 2019). Next, we investigated mitochondrial and lysosomal dysfunction through immunostaining and examination into associated DEGs. Mitotracker CMXRos staining showed an increase in the number of mitochondria per cell, suggesting an accumulation of mitochondria with time in these IPZ-treated cells (Fig 3D,G). Consistent with these observations, we observed a broad phasic dysregulation of the top upregulated and downregulated DEGs associated with mitochondrial function, suggesting that the expression of mitochondrial genes is strongly correlated with the cell-cycle dysfunction in our model (Fig S4). Investigations into lysosomal dysfunction through immunostaining with lysotracker similarly showed an increased accumulation of lysosomes per cell, suggesting a lack of lysosomal turnover in treated cells (Fig 3D,H). Further investigations into the transcriptional regulation of lysosomal DEGs showed time-dependent lysosomal dysregulation, with clusters of upregulated genes at 48H, as well as 96H and 168H (Fig S5). Of these lysosomal genes, we observed an early upregulation of *LAMP3* as early as 12H, which is maintained throughout the time course (Fig S5). However, other lysosomal genes showed a more gradual upregulation in expression, with many of them showing the highest differential expression at 48H which is then either maintained (*CTNS*, *STS*, *HPS1*, *TPP1*) or observed to reutrns to baseline (*CTSF*) (Fig S5).

**Figure 3:**
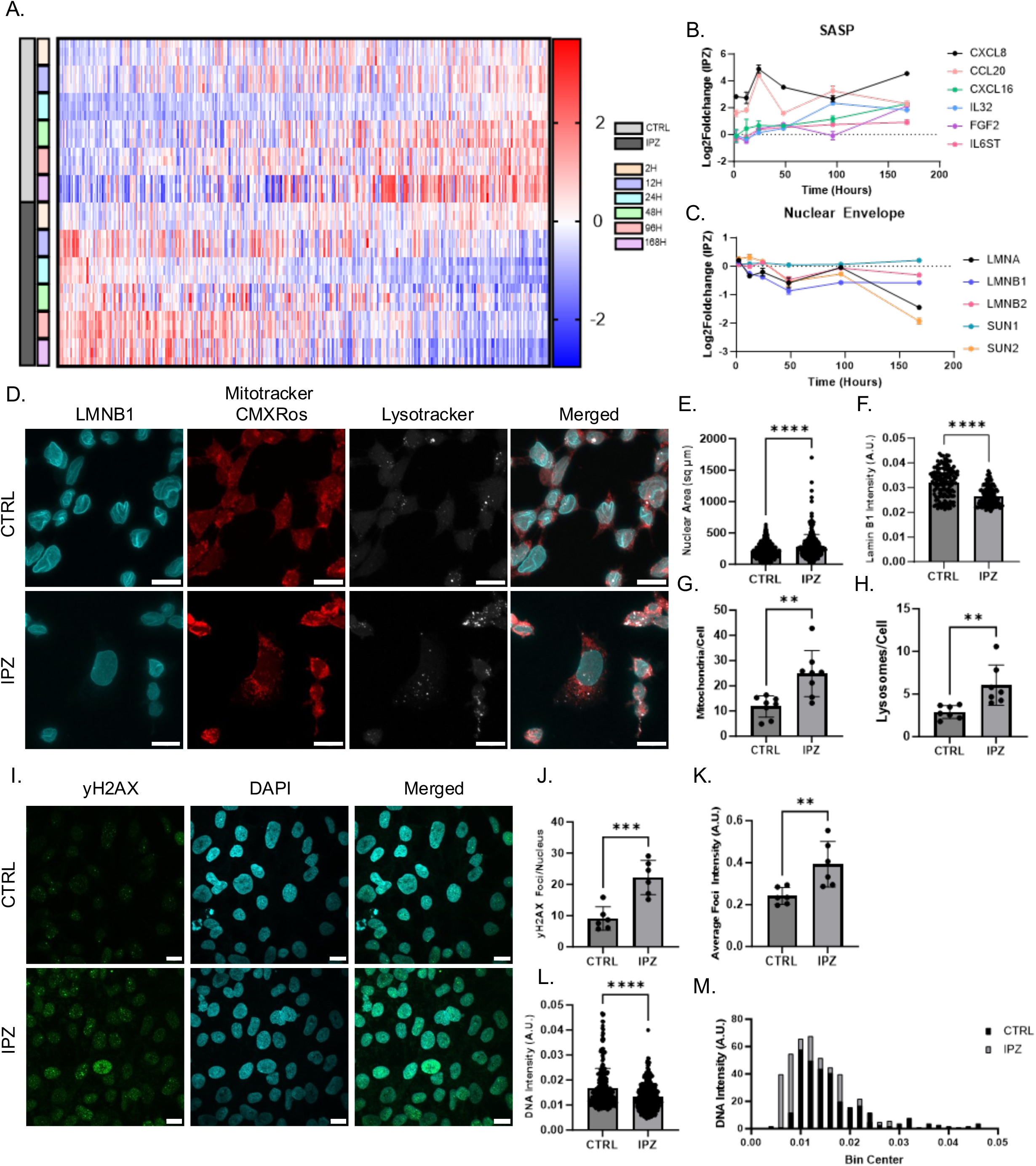
Cell-cycle dysregulation is associated with hallmarks of senescence independent of CKI expression. A) Z-score heatmap of genes associated with senescence and SASP. B-C) Log2FoldChange Expression of SASP (B) and Nuclear Envelope (C) DEGs over the time course of 7-days. D) Immunostaining of lamin B1 (blue) with Mitotracker CMXRos (red) and Lysotracker stains (white) in CTRL and IPZ-treated SK-N-MC cells. E) Nuclear area (μm^2^) of CTRL and IPZ-treated SK-N-MC cells (n=304-379 cells). F) lamin B1 fluorescence intensity from (D) (n=128-134). G-H) Quantification of average mitochondria and lysosomes per cell isolated from Mitotracker CMXRos and Lysotracker stains from (D) (n=8 trials). I) Immunostaining of γH2AX (Ser 139, green) and DAPI (blue) in CTRL and IPZ-treated SK-N-MC cells. J) Quantification of average γH2AX foci per nucleus isolated from (I). K) Average γH2AX fluorescence intensity per foci from (I) (n=6). L) DNA intensity from DAPI staining from isolated from CTRL and IPZ-treated SK-N-MC cells from (I) (n=303-421). All data was analyzed by unpaired two-tailed t-test. (**p<0.01, ***p<0.001, ****p<0.0001). M) Frequency distribution histogram of DNA intensity from (L) for CTRL and IPZ-treated SK-N-MC cells. Scale bars are 10µm.

Lastly, we examined the presence of DNA damage in these IPZ-treated cells at 168H through immunostaining of molecular marker γH2AX (Ser 139). We observed a significant increase in both the number of γH2AX foci per nuclei and the average γH2AX foci intensity with IPZ treatment (Fig 3I-K). We then looked at the distribution of DNA intensity through DAPI staining in these cells to look for DNA replication events through 2n (Bin Center 0.005-0.02+/-0.005) and 4n (Bin Center 0.025-0.040+/-0.005) DNA content populations. Consistent with our observations through FACS (Fig S1), control cells displayed a left shift (increased 2n or G_1_) in DNA content, with a handful of cells displaying a DNA replication event (i.e. 4n DNA content). In contrast, IPZ-treated cells show a wider left skew distribution, with little to no cells in the 4n DNA content range (Fig 3M).

Overall, we demonstrate that cell-cycle dysregulation and arrest contribute to the induction of an immune response in the form of SASP, the downregulation of lamin and nuclear envelope regulators, mitochondrial and lysosomal dysfunction, and ultimately, DNA damage, all of which are classical hallmarks of a senescence-like phenotype.

### *Nemf*^R86S^ MEFs demonstrate G_1_/S cell-cycle arrest and *Stmn2* downregulation

We previously described that *Nemf*^R86S^ mouse embryonic fibroblasts (MEFs) showed an importin-β specific nuclear import defect consistent with our observations in IPZ-treated human neuronal cell lines. This prompted us to investigate whether the R86S mutation was capable of inducing CCD similar to the observed dysregulation we describe in IPZ-treated cells. To test this hypothesis, we investigated the differential expression of E2Fs, cyclins, and CKIs in this R86S MEF model. We observed a slight yet significant downregulation of *E2f1*, as well as an increase in *E2f8*, with insignificant changes in *E2f2/3* transcripts (Fig 4A). We further observed differential expression of cyclins, with *Ccnd1* being downregulated and *Ccne1* being upregulated (Fig 4B). Interestingly, we observed no dysregulation of *Cdkn1a* and *Cdkn2d,* but a large downregulation of the *INK4* locus *Cdkn2a/b* (Fig 4C). This dysregulation of cell-cycle machinery was consistent with the observed dysregulation of key LNCRNAs such as *Neat1*, *Malat1*, *Tug1*, and *Hotairm1*, which have been observed to be dysregulated in neurodegeneration and our IPZ-treated human cells (Fig 4D). Further investigation into protein expression of E2F1 and p16^Ink4a^ show significant downregulation (Fig 4E-G), consistent with the observed transcriptional dysregulation (Fig 4A). We then utilized a semi-quantitative antibody array for proteins expressed in the cell cycle and observed aberrant activation of many key proteins such as variants of phospho-TP53, cyclin E1, and Cdc2, as well as the downregulation of Smad3 and Chk2 (Fig S6).

**Figure 4:**
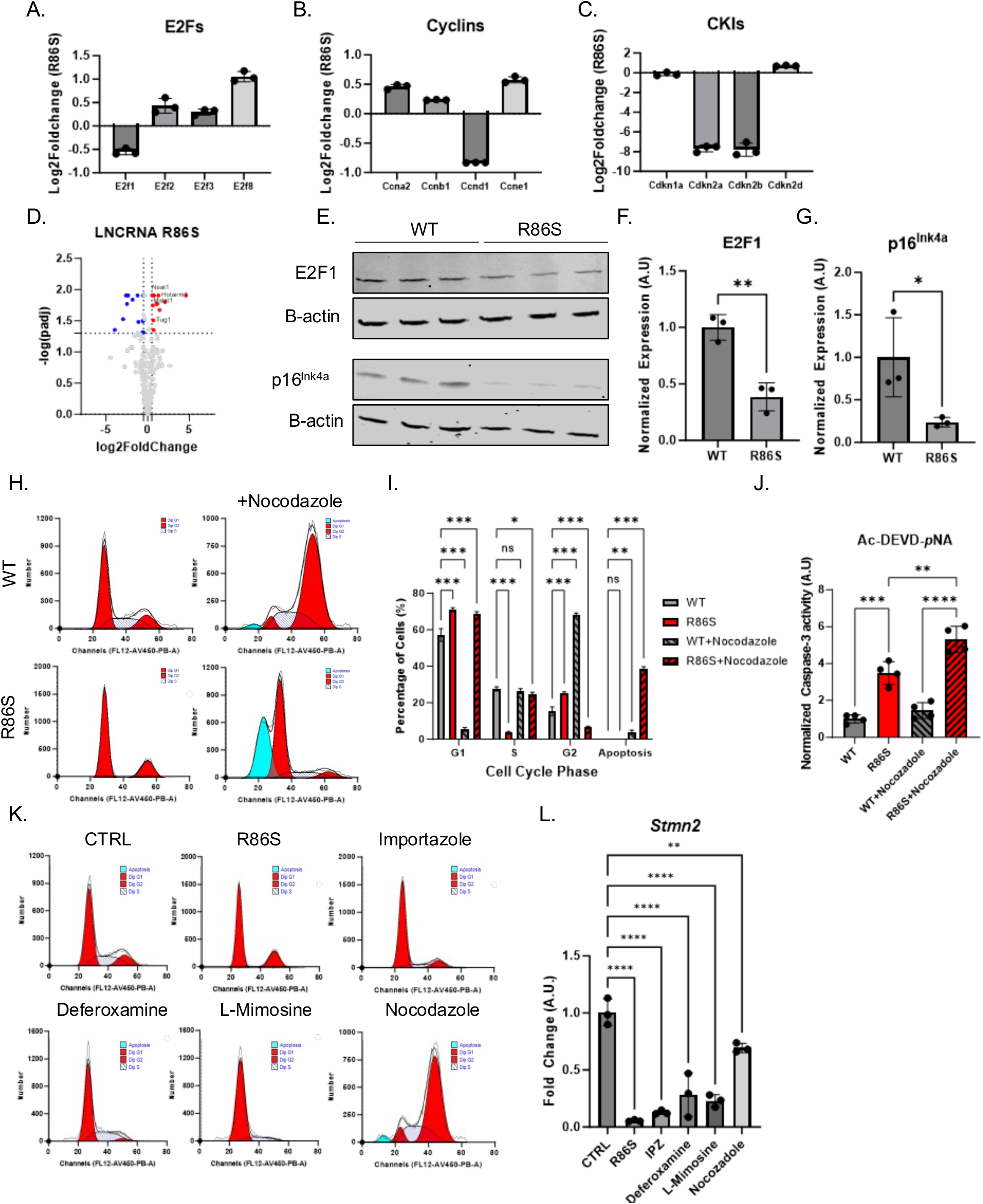
*Nemf*^R86S^ MEFs demonstrate a G_1_/S cell-cycle arrest and *Stmn2* downregulation. A-C) Log2FoldChange Expression from *Nemf*^R86S^ MEFs for E2Fs (A), Cyclins (B), and CKIs (C). D) Volcano plot analysis of Long non-coding RNA DEGs. Red data points indicated upregulated DEGs (log2FC>0.5, padj<0.05) and blue data points indicate downregulated DEGs (log2FC<-0.5, padj<0.05). Gray data points indicate DEGs that do not meet the threshold (-0.5<log2FC<0.5|padj>0.05). E) Western blot analysis of E2F1 and p16^INK4A^ with respective β-actin loading control. F-G) Quantification of protein expression from western blot analysis in (E). Data analyzed by unpaired two-tailed t-test (n=3). H) FACS of DNA content from DAPI staining of WT and *Nemf*^R86S^ MEFs with and without Nocodazole treatment. I) Quantification of the percentage of cells from (H) separated into G_1_, S, G_2_, and apoptosis peaks based on DNA content. Data from (H) analyzed by two-way ANOVA with Šídák’s multiple comparisons test (n=3). J) Colorimetric Caspase-3 activity assay measuring AC-DEVD-pNA cleavage for WT and *Nemf*^R86S^ MEFs with and without Nocodazole treatment normalized to WT (n=4) K) FACS of DNA content from DAPI staining of Control (WT) and R86S MEFs, as well as WT MEFs treated with Importazole, Deferoxamine, L-Mimosine, and Nocodazole. L) Quantitative PCR analysis of *Stmn2* RNA isolated from Control (WT) and R86S MEFs, as well as WT MEFs treated with Importazole, Deferoxamine, L-Mimosine, and Nocodazole (n=3). Data from (J,L) analyzed by ordinary one-way ANOVA with Tukey’s multiple comparison test. (ns p>0.05, * p<0.05, ** p<0.01, *** p<0.001, **** p<0.0001).

Next, we utilized FACS of DAPI-stained WT and *Nemf*^R86S^ MEFs which shows the distribution of DNA content (Fig 4H). WT MEFs showed a G_1_/S/G_2_ distribution of 55/28/17 (%), highlighting that most cells exist in G_1_, with some cells undergoing mitosis by cycling through S/G_2_ phase (Fig 4H). The *Nemf*^R86S^ MEFs showed a 71/4/25 (%) distribution, suggesting that most cells are arrested in G_1_ and G_2_, with few cells undergoing DNA replication in S phase (Fig 4I). To then further confirm if the cells are arrested in G_1_ or G_2_, we induced a G_2_ arrest through nocodazole treatment, which is used to disrupt microtubules, arresting cells in G_2_/M as seen through treatment of WT MEFs (Fig 4H,I). Interestingly, *Nemf*^R86S^ MEFs failed to arrest in G_2_ and remained in G_1_, suggesting that a majority of the cells are arrested in G_1_/S and cannot pass the restriction checkpoint (Fig 4H,I). Concurrently, Nocodazole treatment induced an increase in apoptosis in both WT and *Nemf*^R86S^ MEFs as indicated by fractionated DNA as observed in FACS, with the *Nemf*^R86S^ MEFs showing a significant percentage of cells in this apoptosis peak (Fig 4H,I). This significant increase in apoptosis is in line with an increase in caspase-3 activity as measured through AC-DEVD-pNA cleavage in the *Nemf*^R86S^ MEFs (Fig 4J).

Our previous characterization of *Nemf*^R86S^ MEFs showed that the importin-β specific nuclear import defect was consistent with the dysregulation of key genes in neurodegeneration. Among these dysregulated transcripts, we had previously described *Stmn2* to be significantly downregulated (Plessis-Belair et al., 2024). Thus, we wanted to determine if bypassing the nuclear import block and inducing cell-cycle arrest can similarly result in the dysregulation of *Stmn2*. Treatment of MEFs with IPZ resulted in a G_1_/S arrest as shown through FACS DNA intensity distributions and similarly resulted in a significant downregulation of *Stmn2* (Fig 4K,L). Therefore, we utilized Deferoxamine and L-Mimosine which have been previously shown to arrest cells in the G_1_/S phase (Fukuchi et al., 1997; Park et al., 2012). Deferoxamine is a known iron chelator resulting in iron deprivation resulting in late G_1_ arrest, while L-Mimosine has been shown to inhibit DNA replication resulting in an early S phase arrest (Fukuchi et al., 1997; Park et al., 2012). Transient treatment of WT MEFs with both of these pharmacological agents indeed induced G_1_/S cell cycle arrest as well as a significant downregulation in *Stmn2* transcript levels (Fig 4K,L). Interestingly, cell cycle arrest in G_2_/M utilizing Nocodazole resulted in a slight reduction in *Stmn2* transcript levels, but not to the degree as observed with G_1_/S arrest (Fig 4L).

Overall, these findings suggest that the *Nemf*^R86S^ mutation results in CCD consistent with a G_1_/S cell cycle arrest and that cell cycle arrest in G_1_/S is sufficient to induce *Stmn2* transcript downregulation in a mouse mitotic cell line. These findings highlight an important key feature that CCD may be a consequence of a nuclear import block, yet upstream of the transcriptional dysregulation observed in neurodegeneration.

### Mutant *Nemf*^R86S^ and IPZ-treated primary neuronal cultures demonstrate time-dependent transcriptional dysregulation consistent with cell cycle dysregulation

We isolated primary cortical neurons from WT and *Nemf*^R86S^ mice from the same litter at P0 and cultured the isolated cells for 2 weeks. At this 2-week time point, we treated WT primary neurons with 20µM IPZ for 2 days (2 weeks, 2 days) and 7 days (3 weeks) and simultaneously maintained the respective control (CTRL) and *Nemf*^R86S^ neurons in culture for each respective time point (Fig 5A). At a given time point, we extracted RNA and performed RT-qPCR for target genes (*Stmn2, E2f1, Cdkn1a, Cdkn2a, Meg3, Lmnb1, Cxcl8, Il6*) previously observed to be dysregulated in IPZ-treated SK-N-MC cells (Fig 2) and *Nemf*^R86S^ MEFs (Fig 4). Consistent with our previous study, we observed a significant downregulation of *Stmn2* transcripts in both the R86S and IPZ-treated primary neurons at both 2 days and 7 days (Fig 5B). Interestingly, we observed no significant dysregulation of *E2f1* at 2 days, but an upregulation of *E2f1* in IPZ-treated primary neurons at 7 days (Fig 5C).

**Figure 5:**
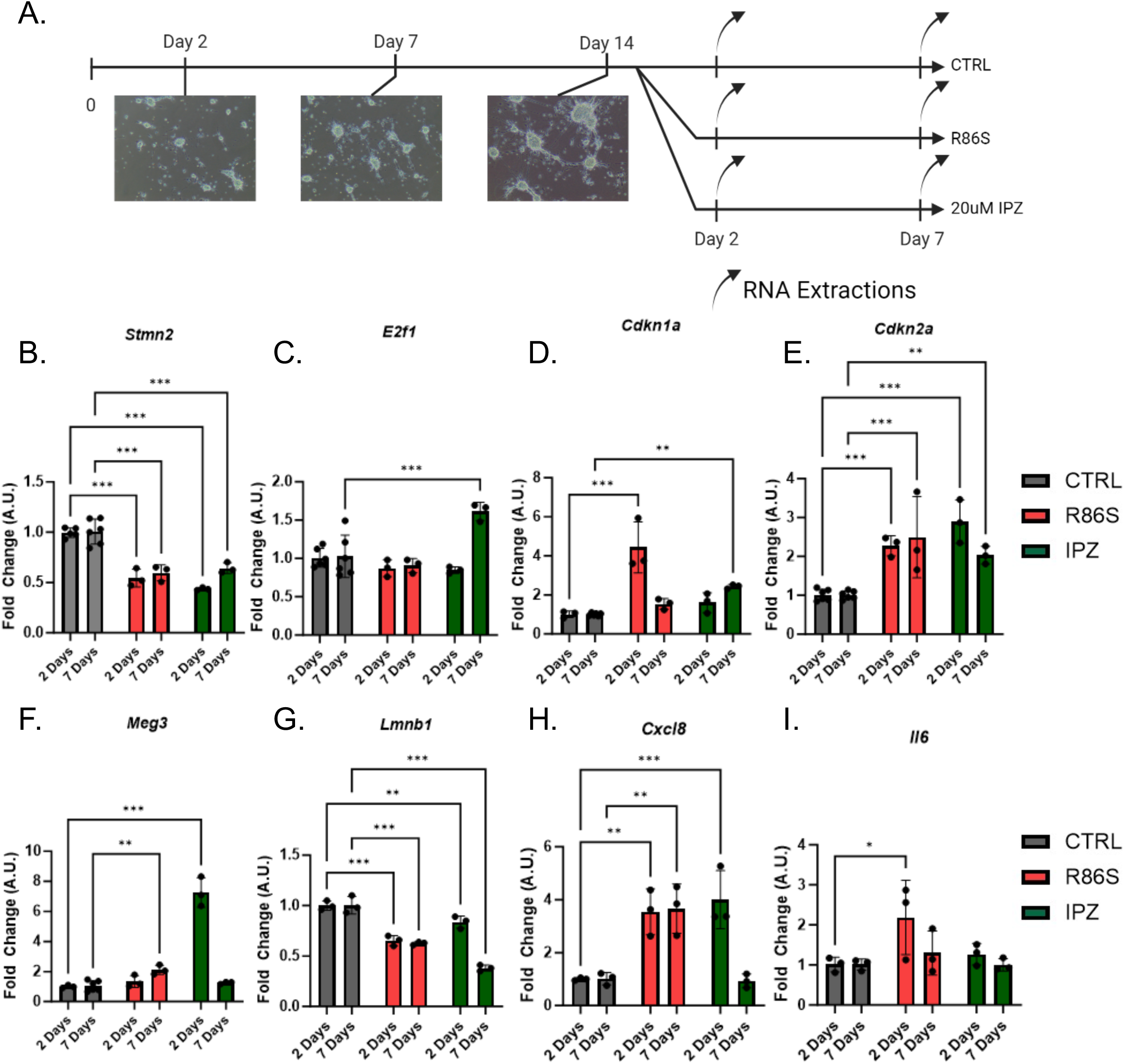
Mutant *Nemf*^R86S^ and IPZ-treated primary neuronal cultures demonstrate time-dependent transcriptional dysregulation consistent with cell-cycle dysregulation. A) Timeline of the experimental set up. Primary cortical neurons were isolated at P0 from the same litter and cultured for 2 weeks. At the 2-week time point, CTRL cells (WT mice) were treated with IPZ (20µM) for either 2 or 7 days. CTRL and *Nemf*^R86S^ (R86S) primary cultures were maintained in parallel with IPZ-treated cells, with RNA extractions at 2 and 7 days. B-I) Quantitative PCR of CTRL, R86S, and IPZ-treated primary cortical neurons of *Stmn2* (B)*, E2f1* (C)*, Cdkn1a* (D)*, Cdkn2a* (E)*, Meg3* (F)*, Lmnb1* (G)*, Cxcl8* (H)*, Il6* (I) at 2 and 7 days. Data from (B-I) analyzed by two-way ANOVA with Šídák’s multiple comparisons test. (n=3) (* p<0.05, ** p<0.01, *** p<0.001).

Next, we investigated the expression of CKI genes *Cdkn1a* and *Cdkn2a,* which have been shown to be upregulated in neurons following aberrant cell-cycle re-entry and neurescence (Hudson et al., 2024). The expression of *Cdkn1a* and *Cdkn2a* is highly variable in neurodegeneration, with the specific upregulation of *Cdkn1a* in neurons being associated with aging and PD (Hudson et al., 2024; Jurk et al., 2012; Riessland et al., 2019) and the upregulation of *Cdkn2a* being more closely associated with AD (Rödel et al., 1996; Vazquez-Villaseñor et al., 2020). We found a significant upregulation of Cdkn1a at 2 days, but not at 7 days in *Nemf^R86S^*(Fig 5D). Concurrently, *Cdkn1a* levels were not significantly different in the IPZ-treated primary neurons at 2 days but were significantly increased at 7 days (Fig 5D). *Cdkn2a* expression in R86S and IPZ-treated primary neurons were found to be significantly upregulated at both 2 days and 7 days, with *Cdkn2a* levels slightly decreasing in IPZ-treated but remaining upregulated relative to control at 7 days (Fig 5E). Lastly, we observed a drastic upregulation in LNCRNA *Meg3* for the IPZ-treated primary neurons at 2 days, which was no longer found to be statistically different at 7 days (Fig 5F). Conversely, *Meg3* showed no significant differences at 2 days but was later found to be significantly upregulated in the R86S primary neurons at 7 days (Fig 5F).

To further validate a senescence-like phenotype we looked at the expression of senescence-associated nuclear envelope gene *Lmnb1* and SASP factors *Cxcl8* and *Il*6 in our primary neuronal cultures. *Lmnb1* was significantly downregulated at 2 and 7 days in the R86S primary neurons consistent with a senescence-like phenotype (Fig 5G). Expression of *Lmnb1* in IPZ-treated primary neurons was slightly downregulated at 2 days and significantly downregulated at 7 days, suggesting a consistent decrease in *Lmnb1* expression with time (Fig 5G). We further observed a significant increase in SASP marker *Cxcl8* expression at 2 days and 7 days in the R86S primary neurons (Fig 5H). However, the expression of *Cxcl8* increased at 2 days IPZ-treated primary neurons but returned to control levels at 7 days (Fig 5H). Expression of *Il6* in the R86S was significantly upregulated with high variation across replicates at 2 days, but insignificant at 7 days (Fig 5I). *Il6* expression was insignificant at both time points in IPZ-treated primary neurons (Fig 5I).

Altogether, the transcriptional dysregulation observed suggests neuronal cell-cycle re-entry through the aberrant expression of CKIs *Cdkn1a* and *Cdkn2a*. This neuronal associated cell-cycle dysregulation is consistent with the downregulation of *Stmn2* and time-specific expression of LNCRNA *Meg3*. Furthermore, the downregulation of senescence-associated *Lmnb1* and upregulation of SASP factors in these post-mitotic neurons suggests senescence-like features following chronic nuclear import defects.

### Differential Expression and localization of CKIs in *Nemf*^R86S^ and IPZ-treated post-mitotic neurons

Based on our observations of transcriptional dysregulation of cell-cycle components and senescence-associated biomarkers, we immunostained CTRL, R86S and IPZ-treated primary neurons at the 7-day (3-week total) time point for CKIs p16^INK4a^ (*Cdkn2a*) and p21^Cip1^ (*Cdkn1a*). The expression of p16^Ink4a^ and p21^Cip1^ under CTRL conditions was relatively low, with faint and diffuse signal in the nuclei of both neuronal and non-neuronal cells (Fig 6A-E). The expression of p16^Ink4a^ in R86S primary cells was found to be uniquely expressed in MAP2+ neuronal cells (Fig 6A). The localization of p16^Ink4a^ was diffuse in the cytoplasm of these R86S primary neurons, with relatively low expression in the nucleus (Fig 6A-C). In contrast, expression of p16^Ink4a^ in IPZ-treated primary neurons was found to be exclusive to the nucleus of MAP2+ cells, with relatively low expression in the cytoplasm (Fig 6A-C). Therefore, the discrepancy between the localization of p16 between the IPZ-treated and R86S primary neurons is not a result defective importin-β nuclear import and suggests activity-dependent localization (Mendaza et al., 2020; Vazquez-Villaseñor et al., 2020). The expression of p21^Cip1^ in R86S and IPZ-treated primary cells were found to be exclusively nuclear, with expression in both neuronal and non-neuronal cells (Fig 6D,E). However, cells expressing the highest levels of p21^Cip1^ were predominantly non-neuronal cells, contrary to the observed p16^Ink4a^ expression (Fig 6E).

**Figure 6:**
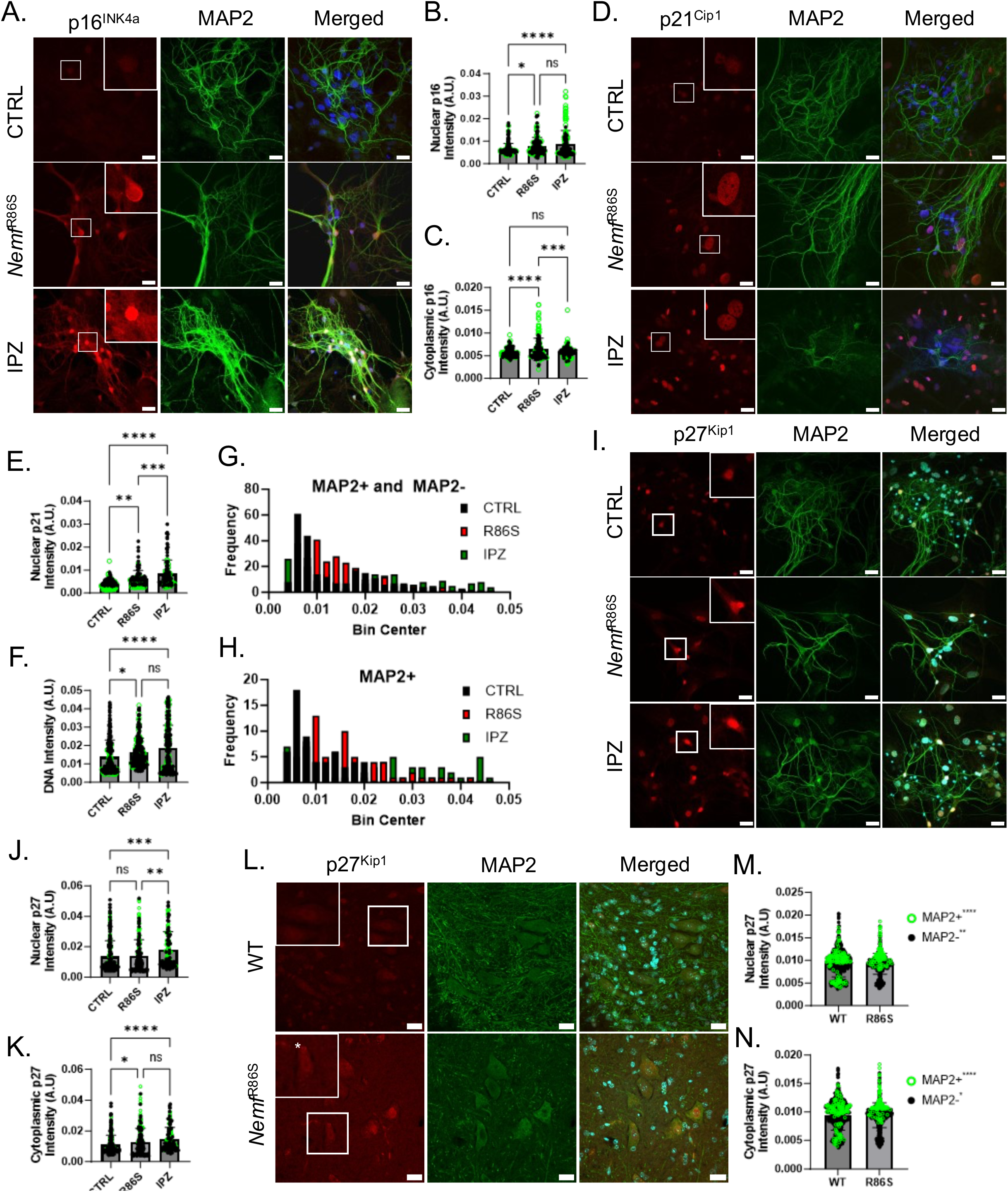
Differential Expression and localization of CKIs in *Nemf*^R86S^ and IPZ-treated post-mitotic neurons. A) Immunostaining of p16 (A, red), MAP2 (green), and DAPI (blue) in CTRL, *Nemf*^R86S^ (R86S), and IPZ-treated primary cortical neuronal cultures. B,C) Nuclear (B) and cytoplasmic (C) p16 intensity from (A) isolated from neuronal (MAP2+, green open circle) and non-neuronal (MAP2-, black closed circle) cells (n=166-185 cells). D) Immunostaining of p21 (red), MAP2 (green), and DAPI (blue) in CTRL, *Nemf*^R86S^ (R86S), and IPZ-treated primary cortical neuronal cultures. E) Nuclear p21 intensity from (D) isolated from neuronal (MAP2+, green open circle) and non-neuronal (MAP2-, black closed circle) cells (n=104-118 cells). F) DNA intensity from DAPI staining isolated from neuronal (MAP2+, green open circle) and non-neuronal (MAP2-, black closed circle) cells (n=225-244 cells). Data from (C-F) analyzed by ordinary one-way ANOVA with Tukey’s multiple comparison test. (ns p>0.05, * p<0.05, ** p<0.01, *** p<0.001, **** p<0.0001). G,H) Frequency distribution histograms from (F) separated into all cells (MAP2+ and MAP2-cells) and MAP2+ cells. I) Immunostaining of p27 (red), MAP2 (green), and DAPI (blue) in CTRL, *Nemf*^R86S^ (R86S), and IPZ-treated primary cortical neuronal cultures. J,K) Nuclear (J) and cytoplasmic (K) p27 intensity from (I) isolated from neuronal (MAP2+, green open circle) and non-neuronal (MAP2-, black closed circle) cells (n=120-266 cells). L) Immunostaining of p27 (red), MAP2 (green), and DAPI (blue) in the ventral horn of WT and *Nemf*^R86S^ spinal cord sections. M,N) Nuclear (M) and cytoplasmic (N) p27 intensity from (L) isolated from neuronal (MAP2+, green open circle) and non-neuronal (MAP2-, black closed circle) cells (n=111-120 MAP2+ cells, 506-536 MAP2-cells). Scale bars are 20µm.

Expression of p16^Ink4a^ and p21^Cip1^ in both R86S and IPZ-treated primary cells highlights cell-cycle re-entry in these neuronal cells consistent with a senescence-like phenotype. We then investigated whether there was also a correlation with an increase in total DNA levels as measured by DNA intensity from DAPI staining (Sigl-Glöckner & Brecht, 2017). We see an overall increase in total DNA staining in the R86S (0.01628 A.U.) and IPZ-treated (0.01843 A.U.) groups relative to control (0.013847 A.U.) which consisted of both MAP2+ and MAP2-cells (Fig 6F). However, DNA intensity distributions for control conditions showed only non-neuronal cells with above-average intensities, suggesting that MAP2+ neuronal cells are not actively cycling (Fig 6F). We then calculated the frequency of cells that fell within a DNA intensity bin range of +/-0.002 (Fig 6G,H). We found that CTLR primary cells correlated with left-skewed distribution, consistent with the predominant population falling in the G_0_/G_1_ range (Fig 6G). Isolating only MAP2+ shows that the majority of these neuronal cells were found within this G_0_/G_1_ range (Fig 6H). In contrast, R86S cells showed a slight rightward shift in DNA intensity in both MAP2+ and MAP2-populations, suggesting that neuronal cells might be undergoing a DNA replication event, or might be further arrested in a late G_1_/S phase (Fig 6G,H). IPZ-treated MAP2+ and MAP2-populations show a wide range of DNA intensities with two distinct populations consistent with a DNA replication event, suggesting that many of these cells including neuronal cells might have undergone polyploidization(Nandakumar et al., 2021) (Fig 6G,H).

Whereas p21^Cip1^ and p16^Ink4a^ have a shared history through their well characterized role in mediating cell cycle progression and promoting cellular senescence, p27^Kip1^, a member of the Cip/Kip family has emerged as a regulator of both cell cycle progression, autophagy, cell cytoskeletal dynamics and apoptosis (Kawauchi et al., 2006; Liang et al., 2007; Polyak et al., 1994; Toyoshima & Hunter, 1994). The functions of p27^Kip1^have been described to be dependent on its subcellular localization with Its ability to regulate the cell cycle being associated with nuclear p27^Kip1^, whereas cytoplasmic p27^Kip1^ enhances cell survival through mediation of autophagy and cytoskeletal dynamics (White et al., 2018). Furthermore, there is strong evidence that the age-related changes in both the expression and localization of p27^Kip1^ plays an integral role promoting quiescence, senescence, and apoptosis in both healthy and aging cells (Liang et al., 2007; White et al., 2018).

Contrary to the expression of p16^Ink4a^ and p21^Cip1^, examinations into the expression and localization of p27^Kip1^ in primary cortical neuronal cultures revealed a diffuse nuclear localization in both neuronal and non-neuronal cells in control conditions (Fig 6I-K). In R86S samples, we observe significant cytoplasmic localization, particularly in MAP2+ cells, but observe no general increase in nuclear expression across cell types (Fig 6I-K). Interestingly, we see both an increase in the nuclear expression and cytoplasmic localization of p27^Kip1^ across conditions, suggesting both an upregulation in protein expression as well as a mis-localization (Fig 6I-K).

We then turned to spinal motor neurons in *Nemf*^R86S^ mice, which were previously described as the source of pathology in this model of neurodegeneration (Martin et al., 2020; Plessis-Belair et al., 2024). Interestingly, these spinal motor neurons do not express p16^Ink4a^ or p21^Cip1^ but were observed to express p27^Kip1^. Immunostaining revealed subpopulations of both neuronal (MAP2+) and non-neuronal (MAP2-) cells which either positively and negatively stained for p27^Kip1^ (Fig 6L-N) In contrast, most spinal motor neurons in *Nemf*^R86S^ mice expressed p27^Kip1^ with increased expression in both the nucleus and the cytoplasm (Fig 6L-N).

Considering the role of CKIs in mediating the cell cycle as well as the role of CKI expression in neurodegeneration, expression of p16^Ink4a^, p21^Cip1^, and p27^Kip1^ in R86S and IPZ-treated neuronal and non-neuronal cells as well as an increase in DNA content in these neurons supports our hypothesis that defective nuclear import mechanisms culminate in cell-cycle re-entry in post-mitotic cells.

## Discussion

When investigating dysregulation of the cell cycle as a potential pathway downstream of nuclear import defects, we must consider the implications of the fluctuating gene expression associated with the baseline mechanisms involved in the cell cycle. In particular, it is important to consider the downstream pathways involved in preparing a cell for mitosis, including but not limited to microtubule reorganization (Hasezawa et al., 1991; Sato & Toda, 2010), DNA synthesis and replication (Sclafani & Holzen, 2007; Stillman, 1996), and nuclear envelope dynamics (Blow & Laskey, 1988; Foisner, 2003). These crucial pathways are often a requirement for proper cycling of mitotic cells, and any perturbations within these pathways would result in cell-cycle arrest, apoptosis, or cancer (Blumenfeld et al., 2017; Chow et al., 2012; Macheret & Halazonetis, 2015; Sallee & Feldman, 2021). However, the occurrence of these cell-cycle associated pathways in post-mitotic neurons, such as the reorganization of microtubules or the nuclear envelope, may have detrimental effects to neuronal health. Therefore, neuronal cell-cycle re-entry events may be unwanted byproducts of integral mechanisms in cellular homeostasis. The evidence provided here for cell-cycle re-entry in both *Nemf*^R86S^ and IPZ-treated primary neurons highlights dysregulation of these crucial cell-cycle associated pathways through the fluctuation of gene expression, downstream of nuclear import defects, culminating in a neurodegenerative phenotype.

Specifically, we focus on the expression of *Stmn2* following cell-cycle arrest at the G_1_/S transition. The activation of the CDK-RB-E2F pathway observed in G_1_, which drives E2F-regulated transcriptional activation, is required for progression through the G_1_/S checkpoint (Johnson et al., 1994; Nevins, 2001; Zhu et al., 2004). It has been previously shown that activation of E2F TFs can upregulate stathmin transcripts through interactions with the stathmin promoter, and thus inhibition of E2F transcriptional activity either through loss of E2F or through inhibition of the CDK-RB-E2F pathway by CKIs can result in decreased stathmin expression (Iancu-Rubin & Atweh, 2005; Polzin et al., 2004). In particular, repression of AP-1 component JUN has been implicated in the reduction of stathmin expression (Kinoshita et al., 2003). Here, we describe the repression of E2F and AP-1 activity downstream of CKI transcriptional downregulation, suggesting that nuclear import defects may initially propagate CCD, independent of CKI inhibition of the CDK-RB-E2F axis. In the case of post-mitotic neuronal cells, we observe aberrant upregulation of CKIs p16^Ink4a^ and p21^Cip1^ associated with cell cycle re-entry, highlighting differences in CKI expression between mitotic and post-mitotic cells. Interestingly, investigations into p27^Kip1^ suggest that the subcellular localization of these CKIs may play an integral role in their function. In particular, p27^Kip1^ has been previously described to inhibit stathmin directly through its cytoplasmic localization, or indirectly through the CDK-RB-E2F axis (Baldassarre et al., 2005; Polzin et al., 2004). Altogether, we suggest that defective nuclear import-induced CCD is a direct upstream pathway resulting in the downregulation of stathmins, specifically *Stmn2,* in NDDs.

Additionally, we describe that these IPZ-treated mitotic cells demonstrate a senescence-like phenotype through the observed expression of the SASP, reduced lamin expression, mitochondrial and lysosomal dysfunction, and DNA damage. The presence of senescence-like hallmarks further suggests that perturbations in cell-cycle regulation alone can drive cellular senescence pathways. Our discrepant observations of CKI expression between mitotic and post-mitotic cells further elucidate a cell-type specific profile of cellular senescence, which are all downstream of cell-cycle dysregulation. Therefore, we suggest a novel and reproducible model to induce cellular senescence *in vitro* by impairing nuclear import and consequently inducing cell-cycle dysregulation.

We have demonstrated here that there is a direct impact on the cell cycle through importin-β-mediated nuclear import inhibition, culminating in cell-cycle dysregulation in mitotic cells as well as inducing cell-cycle re-entry in post-mitotic primary cortical neurons. We further showed the phasic and variable expression of key genes and biomarkers observed to be dysregulated in neurodegeneration, suggesting that cell-cycle re-entry has a strong influence on transcriptional regulation. To this end, we predicted and described repressed transcriptional activity of specific transcriptional regulators in the cell cycle, such as E2Fs and AP-1. This dysregulation in transcriptional activity was further associated with the downregulation of *STMN2*. This pathological cell-cycle re-entry was further recapitulated in the mutant *Nemf*^R86S^ post-mitotic primary cortical neurons which display a significant upregulation of CKIs *Cdkn1a* (p21^Cip1^) and *Cdkn2a* (p16^Ink4a^) both through quantitative PCR and immunostaining, with p16^Ink4a^ expression being neuronal specific and p21^Cip1^ being broadly expressed across cell types. We further show that p27^Kip1^ protein expression is upregulated in both the nucleus and the cytoplasm in our R86S and IPZ-treated primary neurons, as well as in R86S spinal motor neurons. Lastly, we describe in these R86S and IPZ-treated primary neurons a significant downregulation of *STMN2*, and upregulation of the SASP factors *Cxcl8 and Il6* as well as downregulation of *Lmnb1.* Overall, the data suggests that cell-cycle dysregulation is downstream of importin-β nuclear import defects, culminating in cell-cycle re-entry in post-mitotic neurons and models of neurodegeneration.

## Acknowledgments

We wish to thank current and former members of the Sher lab for critical discussions and advice. We wish to thank Dr. Gregory Cox (Jackson Laboratory and University of Maine) for help with providing NEMF mutant mice and with suggestions for research strategies. We thank Dr. Joshua Dubnau for experimental design advice. We thank Wendy Ackmentin for her invaluable help on all aspects of running lab facilities.

## Conflict of Interest statement

The authors declare no competing interests or conflicts of interest.

## Funding statement

Funding provided by startup funds to RBS from Stony Brook University, from the WaterWheel Foundation to RBS, and by National Institutes of Health R01AG079898 to RBS

## Authors’ contributions

**Conceptualization:** Roger Sher, Jonathan Plessis-Belair. Markus Riessland, Taylor Russo

**Formal analysis:** Roger Sher, Jonathan Plessis-Belair

**Funding acquisition:** Roger Sher

**Investigation:** Roger Sher, Jonathan Plessis-Belair

**Methodology:** Roger Sher, Jonathan Plessis-Belair,

**Project administration:** Roger Sher

**Resources:** Roger Sher

**Validation:** Roger Sher, Jonathan Plessis-Belair

**Visualization:** Roger Sher, Jonathan Plessis-Belair

**Writing– original draft:** Roger Sher, Jonathan Plessis-Belair

**Writing– review & editing:** Roger Sher, Jonathan Plessis-Belair, Markus Riessland, Taylor Russo

## Notes

### Competing Interest Statement

The authors have declared no competing interest.

