## Supplementary material for "Nuclear Import Defects Drive Cell Cycle Dysregulation in Neurodegeneration": Supplmentary files

**Supplementary Figures**

### Figure S1

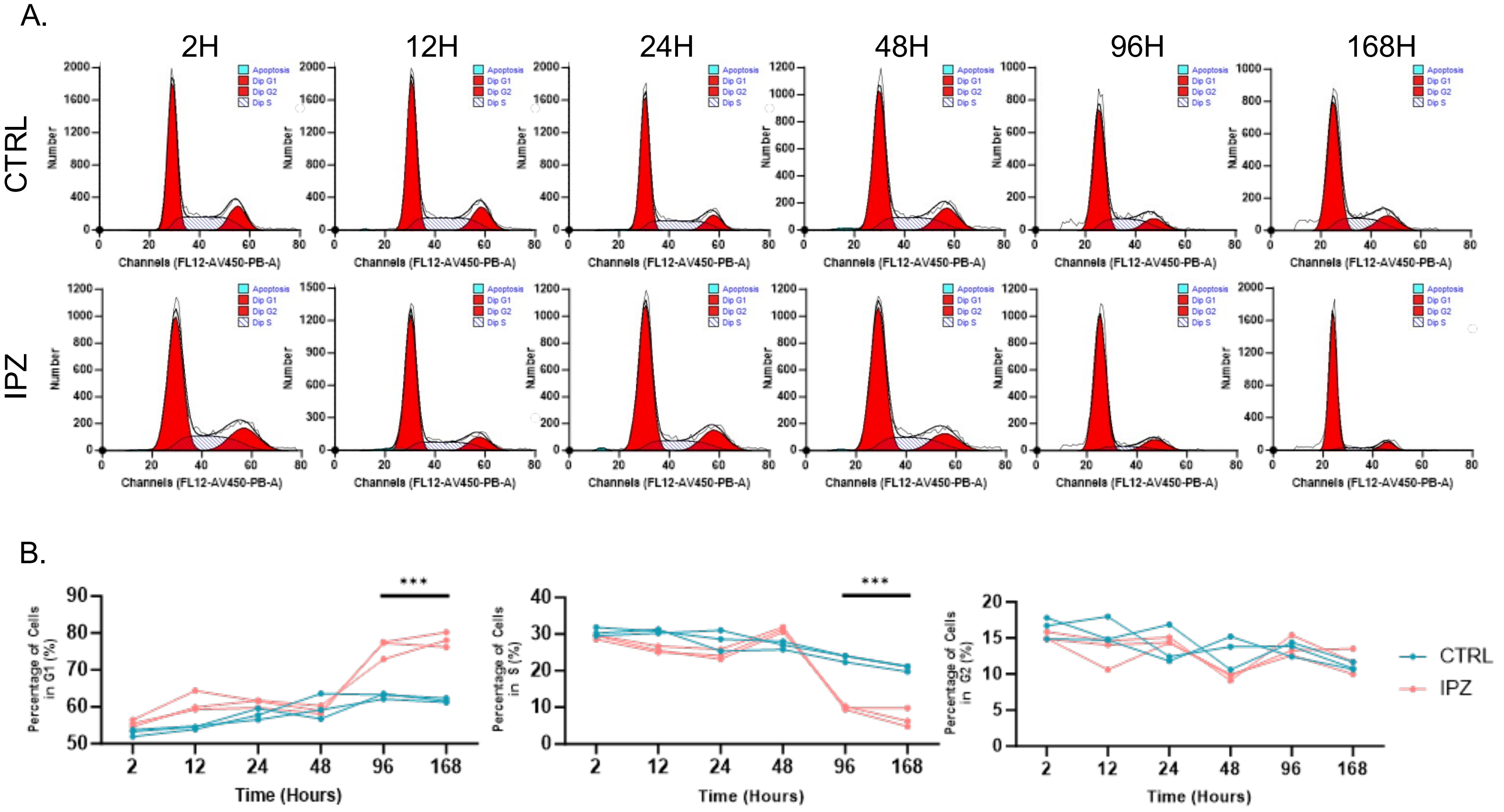


**Fig S1: IPZ-treated mitotic neuronal cell lines display a $G_1$/S cell-cycle arrest through DNA distribution.** A) FACS of DNA content from DAPI staining of Control and IPZ-treated SK-N-MC cells for 2, 12, 24, 48, 96, and 168 hours. B) Percentage of cells from (A) found in the $G_1$, S, and $G_2$ cell cycle phase across each time point.

### Figure S2

A.

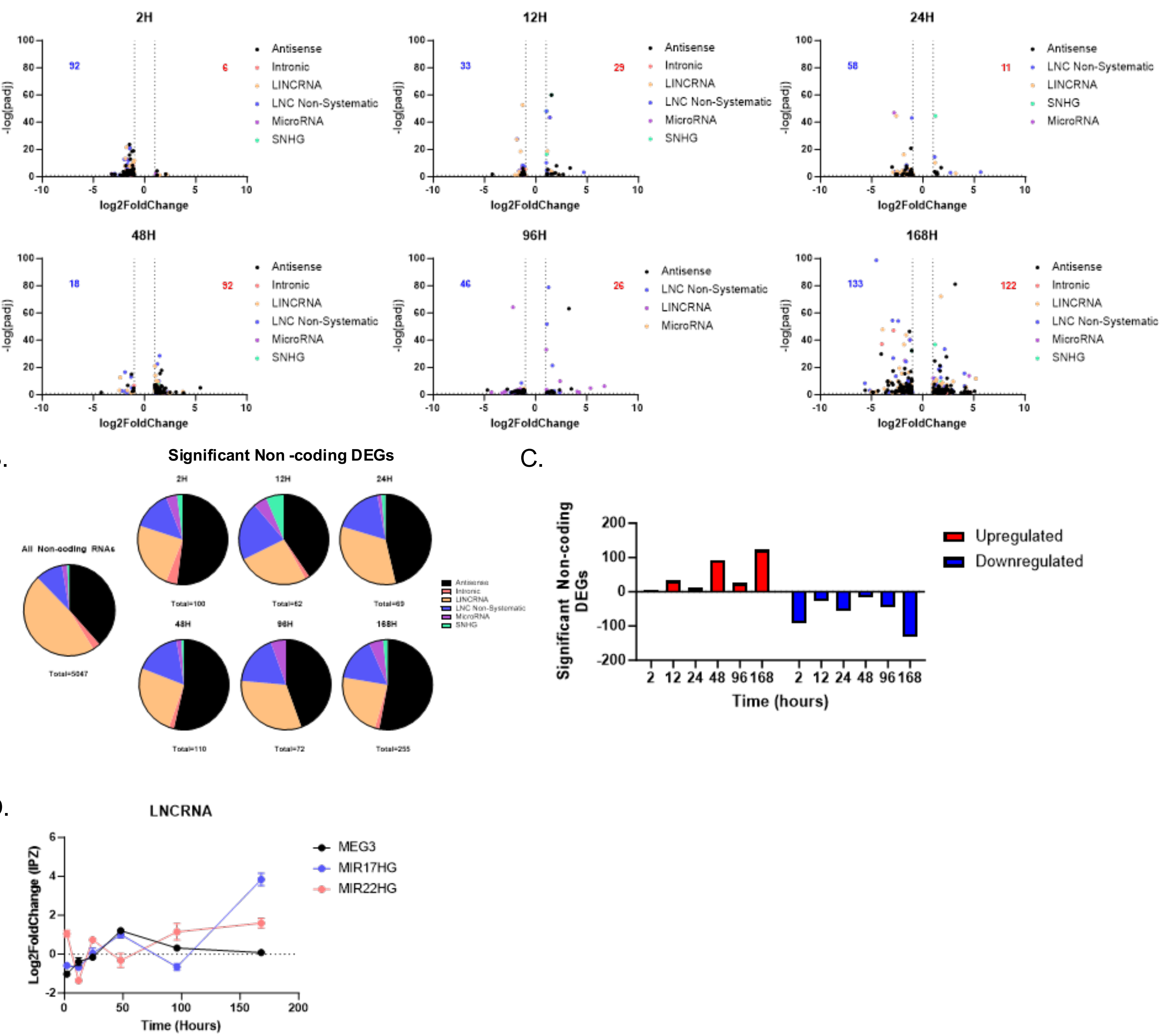


**Fig S2: Dysregulation of LNCRNA DEGs follows cell-cycle associated phasic expression.** A) Volcano plot analysis showing only significant LNCRNA DEGs at 2, 12, 24, 48, 96, and 168 hours separated into Antisense, Intronic, LINCRNA, LNC Non-

Systematic, MicroRNA, and SNHG (padj<0.05, log2FC<-1|log2FC>1). B) Distribution (%) of significant non-Coding DEGS from (A). C) Quantification of upregulated and downregulated significant non-coding DEGs over the 7-day time-course. D) Log2FoldChange Expression of LNCRNA *MEG3*, *MIR17HG*, and *MIR22HG* over the 7-day time course.

### Figure S3

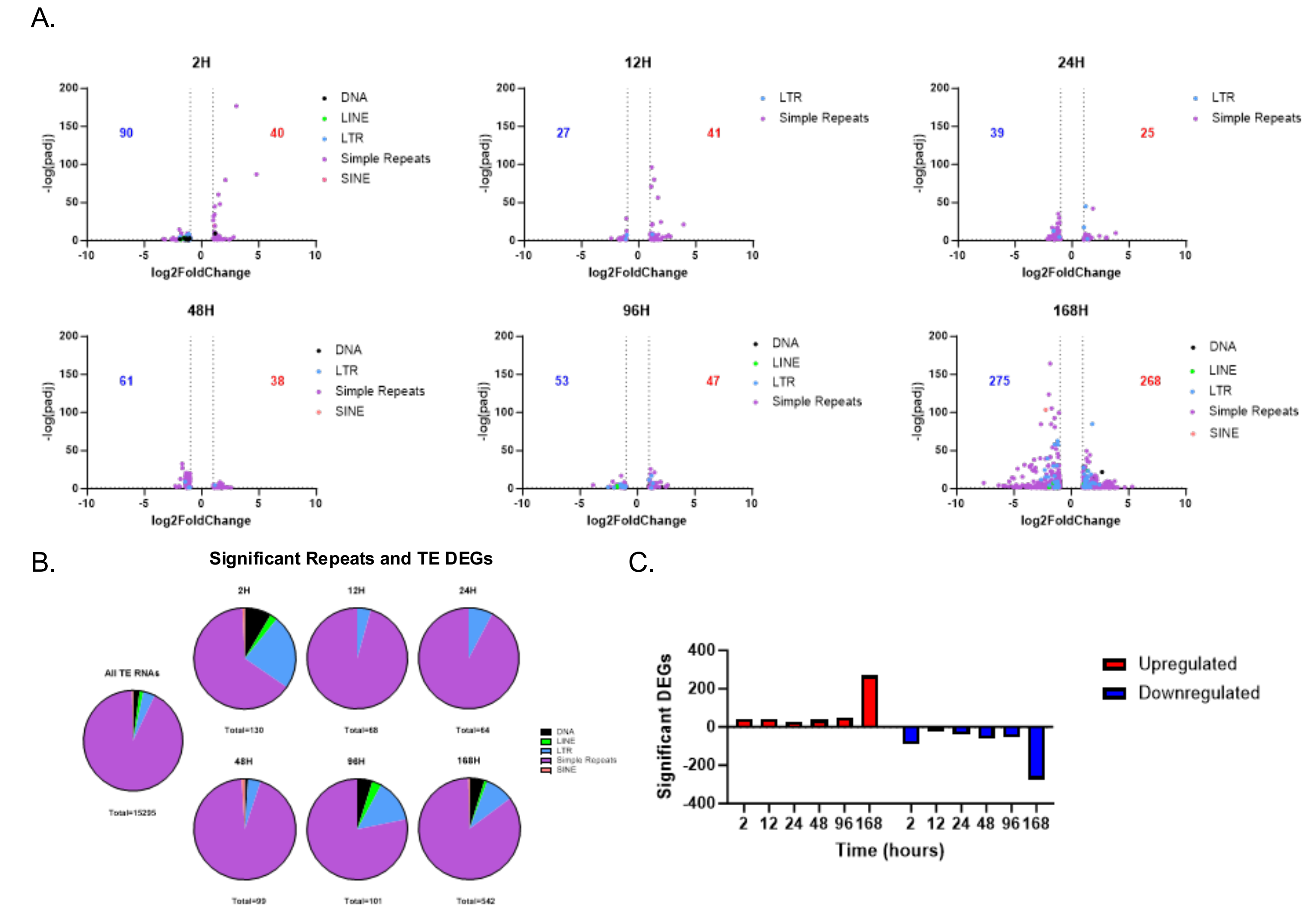


**Fig S3: Dysregulation of Simple Repeats and Transposable Elements occurs late following IPZ treatment.** A) Volcano plot analysis showing only significant Repeat and TE DEGs at 2, 12, 24, 48, 96, and 168 hours separated into DNA Transposons (DNA), LINE, LTR, Sime Repeats, and SINE (padj<0.05, log2FC<-1|log2FC>1). B) Distribution (%) of significant Repeat and TE DEGS from (A). C) Quantification of upregulated and downregulated significant Repeat and TE DEGs over the 7-day time-course.

### Figure S4

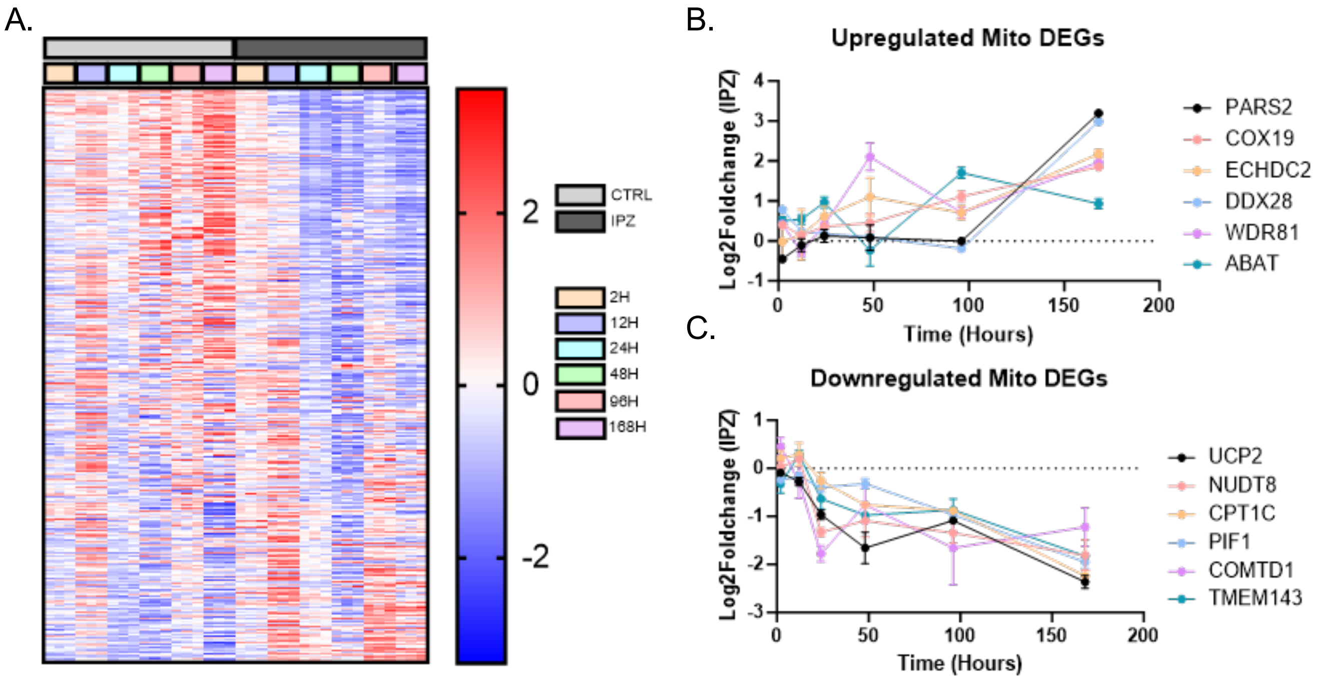


**Fig S4: Upregulation of lysosomal-associated DEGs with IPZ treatment.** A) Z-score heatmap of Mitochondrial DEGs. B-C) Log2FoldChange Expression of Upregulated (B) and Downregulated (C) Mitochondrial DEGs over the time course of 7-days.

#### Figure S5

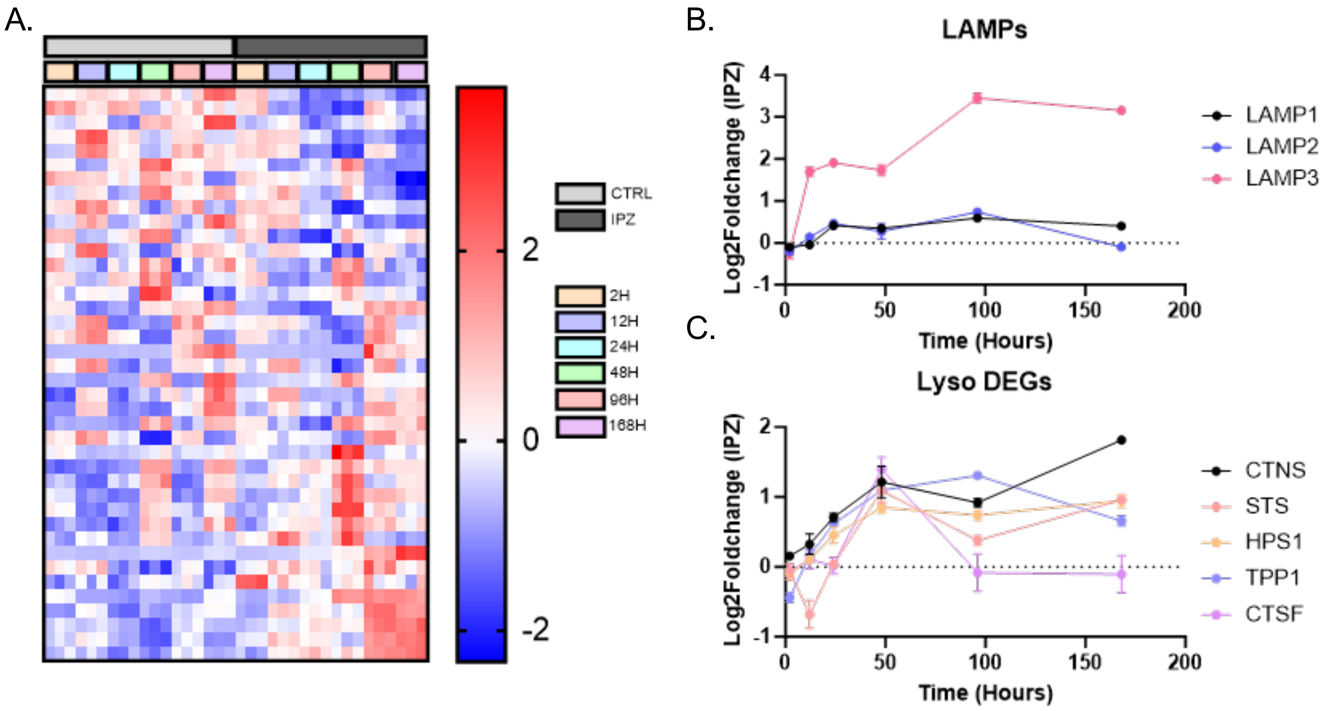


**Fig S5. Cell-cycle dependent dysregulation of mitochondrial-associated DEGs.** A) Z-score heatmap of Lysosomal DEGs. B-C) Log2FoldChange Expression of LAMPs (B) and other Lysosomal (C) DEGs over the time course of 7-days.

**Figure S6**

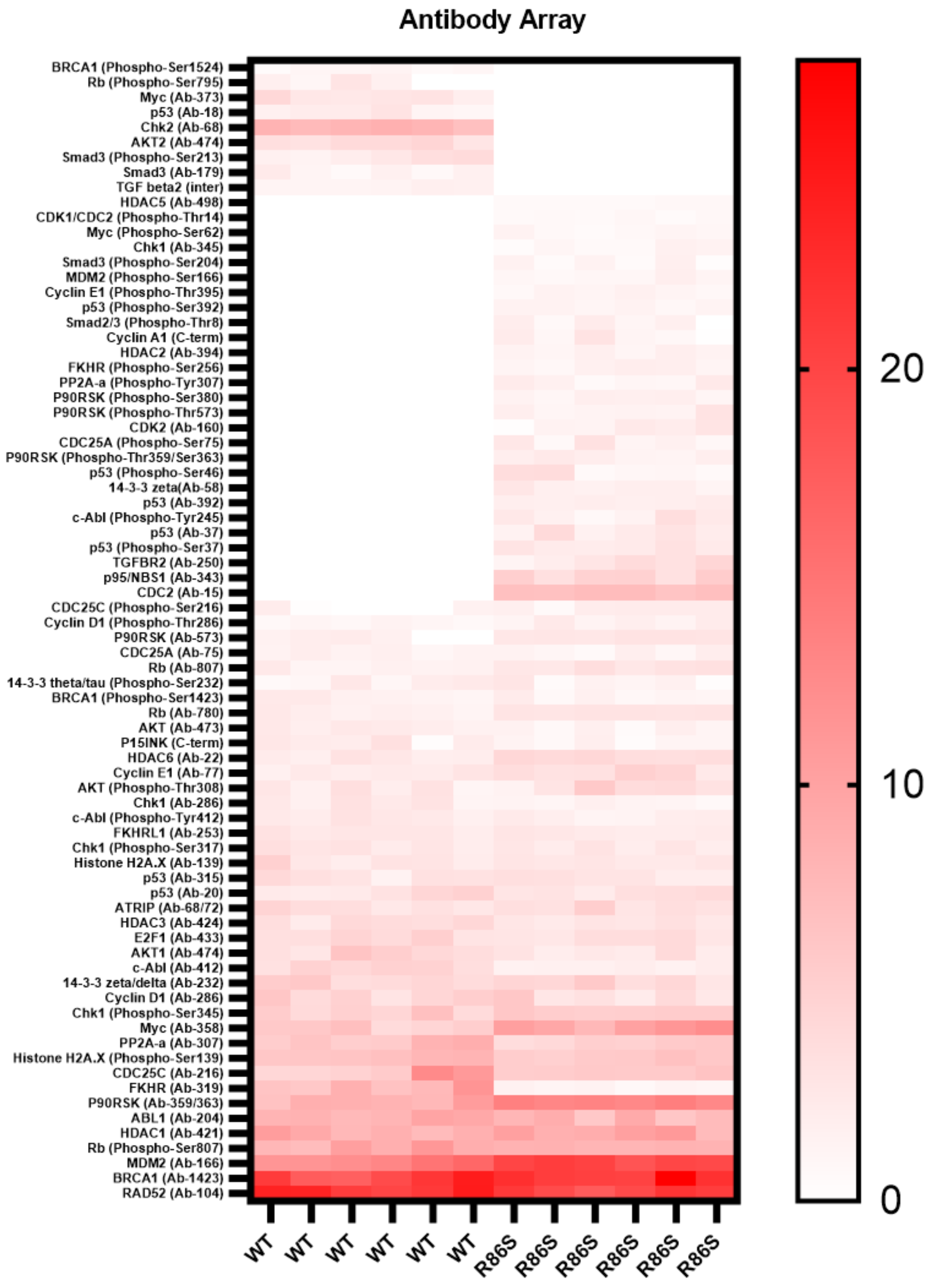


**Fig S6: Phosphorylated Cell-Cycle Protein Expression Antibody Array.** Heatmap of the expression of each detected protein normalized to a beta-actin loading control.

| Cell Culture | Company | Cat # |
|---|---|---|
| Dulbecoo's Modified Eagle's Medium | Corning | 10-017-CV |
| Eagle's Minimum Essential Medium | ATCC | 30-2003 |
| FBS 1X | Gibco | 26140079 |
| Penstrep 100X | Gibco | 15140122 |
| Glutamax 100X | Gibco | 35050061 |
| Non-essential Amino-Acids 100X | Gibco | 11140050 |
| Trypsin-EDTA (0.05%) | Gibco | 25300062 |
| Hibernate-A | Gibco | A1247501 |
| Neurobasal-A | Gibco | 10888022 |
| B27 | Gibco | 17504044 |
| **Antibody** | **Company** | **Cat #** |
| MAP2 | AVES | 4H5 |
| p16INK4a | Invitrogen | PA5-20379 |
| p21 | Invitrogen | MA5-31479 |
| Lamin B1 | Abcam | Ab16048 |
| yH2AX (Ser139) | Cell Signaling | 20E3 |
| Alexa Fluor 594 g@ms IgG2a | Invitrogen | A21135 |
| Alexa Fluor 488 g@Rb Ig | Invitrogen | A11034 |
| Alexa Fluor 488 g@Chk IgY | Invitrogen | A11039 |
| Alexa Fluor 594 g@Chk IgY | Invitrogen | A11042 |
| Alexa Fluor 594 g@ms IgG1 | Invitrogen | A21125 |
| Alexa Fluor 594 g@Rb Ig | Invitrogen | A11037 |
| Alexa Fluor 594 g@ms IgG2b | Invitrogen | A21141 |
| Alexa Fluor 405 g@Rb Ig | Invitrogen | A31556 |
| IRDye 800CW Goat anti-Mouse IgG1 | Licor | 926-32350 |
| IRDye 800CW Goat anti-Rabbit IgG | Licor | 926-32211 |

**Table S1: List of Cell Culture Reagents and Antibodies.**

| Primer | Forward | Reverse |
|---|---|---|
| *Gapdh (mouse)* | GGCAAATTCAACGGCACAGT | GGGTCTCGCTCCTGGAAGAT |
| *Stmn2 (mouse)* | TGTCACTGATCTGCTCCTGC | TGGGAGATGGTGGCTTCAAG |
| *Gapdh (human)* | GTCTCCTCTGACTTCAACAGCG | ACCACCCTGTTGCTGTAGCCAA |
| *Actb (mouse)* | CATTGCTGACAGGATGCAGAAGG | TGCTGGAAGGTGGACAGTGAGG |
| *Cdkn1a (mouse)* | TCGCTGTCTTGCACTCTGGTGT | CCAATCTGCGCTTGGAGTGATAG |
| *Cdkn2a (mouse)* | TGTTGAGGCTAGAGAGGATCTTG | CGAATCTGCACCGTAGTTGAGC |
| *Cxcl8 (mouse)* | CCTTTCCACCCCAAATTTAT | AAACTTCTCCACAACCCTCTG |
| *Il6 (mouse)* | TACCACTTCACAAGTCGGAGGC | CTGCAAGTGCATCATCGTTGTTC |
| *E2f1 (mouse)* | GGATCTGGAGACTGACCATCAG | GGTTTCATAGCGTGACTTCTCCC |

**Table S2: List of Primers.**
